# Distributed control of motor circuits for backward walking in *Drosophila*

**DOI:** 10.1101/2020.07.11.198663

**Authors:** Kai Feng, Rajyashree Sen, Ryo Minegishi, Michael Dübbert, Till Bockemühl, Ansgar Büschges, Barry J. Dickson

## Abstract

How do descending inputs from the brain control leg motor circuits to change the way an animal walks? Conceptually, descending neurons are thought to function either as command-type neurons, in which a single type of descending neuron exerts a high-level control to elicit a coordinated change in motor output, or through a more distributed population coding mechanism, whereby a group of neurons, each with local effects, act in combination to elicit a global motor response. The *Drosophila* Moonwalker Descending Neurons (MDNs), which alter leg motor circuit dynamics so that the fly walks backwards, exemplify the command-type mechanism. Here, we identify several dozen MDN target neurons within the leg motor circuits, and show that two of them mediate distinct and highly-specific changes in leg muscle activity during backward walking: LIN156 neurons provide the hindleg power stroke during stance phase; LIN128 neurons lift the legs at the end of stance to initiate swing. Through these two effector neurons, MDN directly controls both the stance and swing phases of the backward stepping cycle. MDN exerts these changes only upon the hindlegs; the fore-and midlegs follow passively through ground contact. These findings suggest that command-type descending neurons can also operate through the distributed control of local motor circuits.

---

The simple act of walking appears effortless, yet involves the exquisitely timed and scaled activation of multiple muscles in each leg. The motor circuits that produce these muscle movements are controlled by descending inputs from the brain and sensory feedback from the legs^1,2^. The descending inputs modulate the overall locomotor pattern, effecting changes in direction or pace in accordance with the animal’s behavioral goals^3,4^. Sensory feedback, including both external and proprioceptive signals, coordinate progression through the stepping cycle and make the fine adjustments needed to adapt to a varying terrain^5^. While this overall organization is now well established, we still have only very limited understanding of how such descending and peripheral inputs control locomotor circuits. Determining how descending and peripheral inputs control motor circuits for walking thus remains an important open question in basic neuroscience, with the potential to inform approaches in human therapy^6^ and robotics^7^.

One of the most striking examples of the descending control of leg motor circuits is the transition from forward to backward walking. Virtually all animals are capable of walking backwards, albeit with varying degrees of elegance. The most obvious change in leg kinematics in backward walking is the phase switch in leg movements – from the leg moving forward in swing phase and backward in stance phase, to forward in stance phase and backward in swing phase. Yet backward walking involves much more than just a simple phase switch, as the amplitude and speed of movement at each joint must also be altered for smooth backward propulsion^8,9^. Additionally, for animals with multiple pairs of legs, the hindlegs become the leading legs in backward walking, and thereby take over the function of probing the oncoming terrain. The transition to backward walking thus provides an excellent paradigm to explore the more general question of how descending inputs from the brain make the myriad adjustments in motor circuits necessary to effect a smooth and coordinated change in walking direction.

Here we begin to investigate the descending control of locomotor circuits for backward walking in *Drosophila*. Flies, like many other animals, retreat if they perceive a threat or obstacle in the path ahead. These sensory cues are believed to act primarily through a single class of descending neurons, called the Moonwalker Descending Neurons (MDNs), which act as command-type neurons for backward walking^10^. Flies with genetically silenced MDNs are unable to walk backwards when they encounter a physical obstacle, whereas freely roaming flies walk backwards if the MDNs are artificially activated^10^. MDNs receive input from neurons conveying mechanosensory cues^11^ (representing a possible obstacle) and from visual neurons that detect a looming stimulus^12^ (a potential threat). The MDNs are also present in larvae, where they mediate backward crawling^13^. Presumably, MDNs control very different motor circuits in the larvae and adult, as the former produce peristaltic waves of the body wall musculature whereas the latter generate the stepping action of legs. How the MDNs couple to these distinct motor circuits is unknown.

By studying leg joint kinematics, we found that the adult MDNs drive coordinated changes in stepping patterns across all three pairs of legs, with the strongest impact on the hindlegs. By combining anatomical and functional approaches, we identify several dozen morphologically distinct candidate MDN target cell types in the ventral nerve cord (VNC). Consistent with our conclusions from the analysis of joint kinematics, the majority of these neurons are located in the third (metathoracic, or T3) segment. Using cell-type specific genetic drivers, we confirmed that many of these cell types are indeed required for backward walking, and analyzed two of them in detail. The LIN156 neurons – specific to the T3 segment – mediate tibia flexion during the power stroke of backward walking. The LIN128 neurons – present in all segments but activated by MDN most strongly in T3 – lift the hindlegs to initiate swing phase. These two MDN-effector neurons thus mediate two critical hindleg movements in backward walking: the power stroke during stance phase (LIN156) and elevation at the start of swing phase (LIN128).

Models for the descending control of motor circuits have generally emphasized either of two extreme scenarios: a command-type of control, in which a single type of descending neuron, like MDN, effects a coordinated set of changes of many muscle movements, versus a more distributed, population-type of control, in which a single type of descending neuron modulates a limited set of motor outputs, with many such descending inputs working in combination to produce a coordinated pattern of muscle movements. Intuitively, command-type descending neurons might be expected to exert a relatively high-level, centralized control on motor circuits, whereas population-type descending neurons might impinge at a lower level, close to the specific motor outputs they control. Our analysis of MDN outputs suggests, however, that command-type neurons can also act in a highly distributed and localized manner: We propose that MDN acts as a command-type neuron because, collectively, its distributed outputs effect a state switch in motor circuit network dynamics – from a resting or forward walking state to a backward walking state.

## Results

### Joint kinematics suggest that hindlegs dominate MDN-triggered backward walking

Before embarking on a detailed investigation of how MDN couples to leg motor circuits, we first sought to better understand the impact of MDN activation on leg movements. Previous studies of MDN-triggered backward walking have examined only the translocation of the body^10-12^ or the pattern of footfalls^10^. It remains to be determined how MDN alters leg joint kinematics, a more direct and higher-dimensional readout of motor circuit activity. To this end, we acquired high-speed (200 fps) videos of tethered flies walking on a suspended ball^14^, and used the DeepLabCut software^15^ to train a neural network to automatically detect the positions of each leg joint (Fig. 1a and Supplementary Video 1). Each of the 6 legs has 3 articulated joints. From proximal to distal, these are the coxa-trochanter, femur-tibia, and tibia-tarsus joints (in *Drosophila*, the trochanter and femur are fused into a single leg segment; Fig. 1b). These joints, as well as the proximal and distal tips of the leg, were located for all three legs on one side of the body in each video frame (Fig. 1a,b). We analysed videos of both spontaneous forward walking and MDN-triggered backward walking (Fig. 1b-h).

**Fig. 1.**
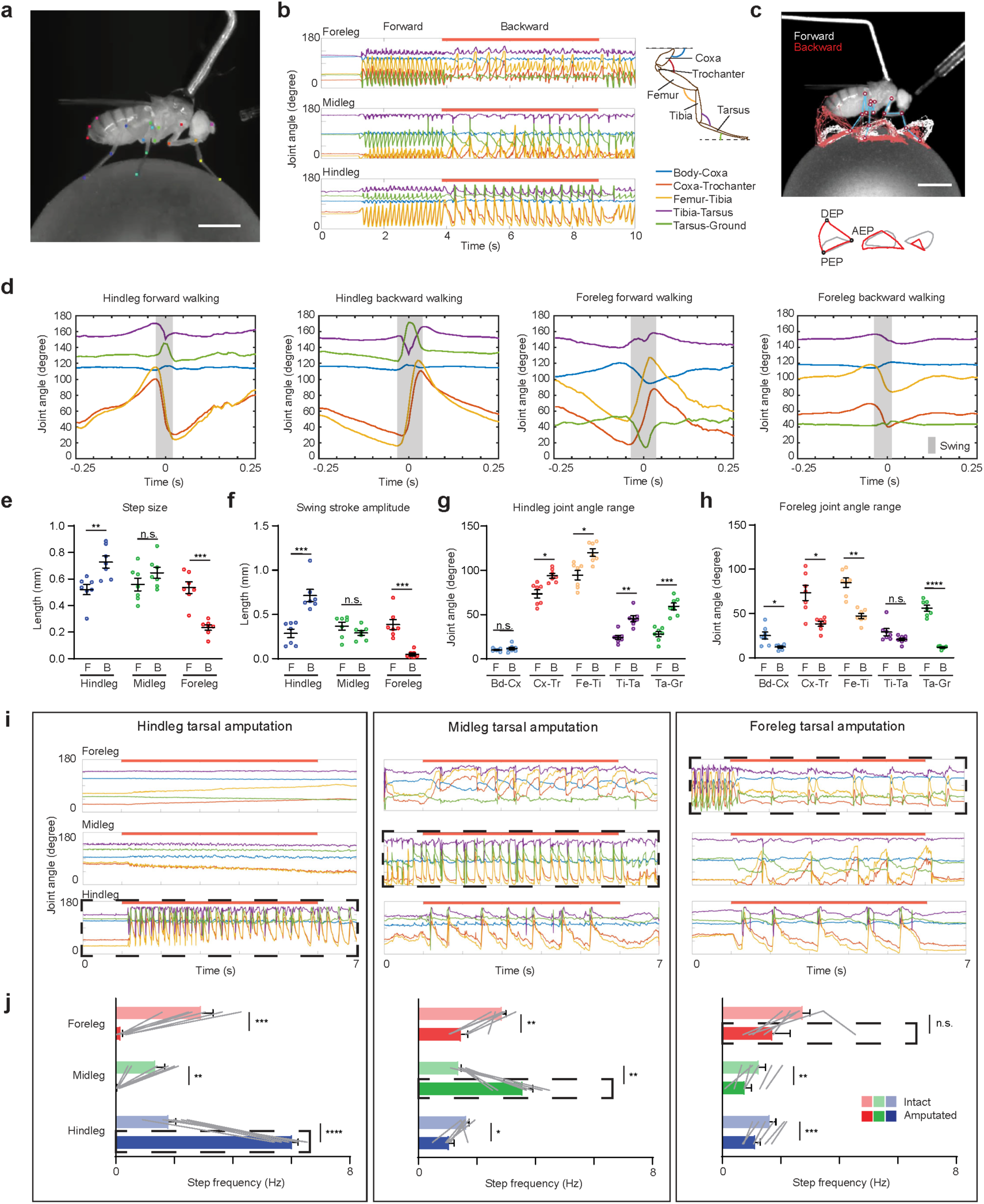
Joint kinematics during forward and backward walking. a, An example frame from high-speed videos of fly walking on a 6-mm diameter ball. Colored dots indicate anatomical landmarks annotated by an artificial neural network trained on DeepLabCut. b, Right, schematic of a *Drosophila* leg showing each leg segment and joint, and color codes used for each joint in all Figures. Left, joint angle tracking from an episode of an *MDN>CsChrimson* fly walking on ball. Red bars indicate the red light stimulus. c, Top, a frame from the Supplementary Video tracked in (b), overlaid with the trajectory of tarsal tips of all three legs during forward walking (white) and backward walking (red). Bottom, average trajectories of tarsal tips during forward walking (grey) and backward walking (red). Anterior (AEP), posterior (PEP) and dorsal (DEP) extreme positions for the hindleg are indicated. See also Supplementary Video 1. d, Joint angles from averaged steps of hindleg and foreleg during forward walking and backward walking. Grey shading represents swing phase, as inferred from the averaged tarsal tip trajectories. *N* = 7 flies; *n* = 156, 75, 156, or 142 steps (from left to right). e, Average step size for each fly during forward and backward walking, defined as the distance between AEP and PEP. F, forward walking; B, backward walking. Bars indicate mean ± s.e.m. of per fly averages. f, Average swing stroke amplitude, defined as the distance between DEP and the line segment connecting AEP and PEP. g, Average foreleg joint angle range (maximum – minimum angle). h, Average hindleg joint angle range. i, Representative joint angle tracking of flies with fore-, mid-, or hindlegs bilaterally amputated at the tarsus. Dashed boxes indicate the amputated leg. See also Supplementary Video 2. j, Stepping frequency for each leg, before and after amputation, shown as mean ± s.e.m.. *n* = 6 flies for each group. Grey lines indicate paired trial averages for the same fly, before and after amputation. ****, *P*<0.0001; ***, *P*<0.001; **, *P*<0.01; *, *P*<0.05; n.s., *P*≥0.05; paired-t-tests, two-tailed *P* values. Scale Bars: 1 mm.

The trajectories of the tarsal tips differed considerably during bouts of forward and backward walking by the same fly (Fig. 1c and Supplementary Video 1). The forelegs stepped further and higher when walking forwards than when walking backwards; the hindlegs took longer and higher strides when walking backwards (Fig. 1c-f). The fore- and hindlegs are generally oriented parallel to the fly’s body axis, and so the lateral placement of our video camera provides the best perspective for tracking joint angles in these legs. We found that the two joints with the greatest range of motion in both forward and backward walking were the coxa-trochanter and femur-tibia joints (Fig. 1b,d,g,h). The flexion and extension of these two joints occurred synchronously for each leg, but in opposite stepping phases for forelegs versus hindlegs (Fig. 1d). During forward walking, the foreleg coxa-trochanter and femur-tibia joints were flexed in stance phase and extended in swing phase; the hindleg joints were flexed in swing and extended in stance. The phase coupling of flexion and extension was reversed during MDN-triggered backward walking. Thus, the forelegs appear to pull the body forward during forward walking, with the hindlegs either pushing or passively extending. For backward walking, the hindlegs pull, while the forelegs either push or passively extend. Regardless of which direction the fly walks, the leading legs take the higher steps, which may facilitate walking on uneven terrain or over obstacles.

To assess whether the hindlegs indeed provide the greatest propulsive force during backward walking, we next performed a series of amputation experiments. In these experiments, we first monitored leg joint kinematics in an intact fly upon MDN activation, then bilaterally amputated the tarsal tips from one pair of legs before again activating MDN and videotaping leg movements. Tarsal amputation prevents the leg from gaining any traction on the ground but preserves all the leg joints. We found that the fore- and midleg amputees were still able to walk backwards, whereas the hindleg amputees could not (Fig. 1i,j and Supplementary Video 2). Thus, it is primarily the stance phase power stroke of the hindleg that drives the body backwards during MDN-triggered backward walking. Interestingly, in all cases, the amputated legs continued to oscillate, suggesting that their stepping motion is actively driven by central motor circuits and not merely a passive consequence of their surface traction (Fig. 1i,j and Supplementary Video 2). We also noticed that, particularly for the hindlegs, but to a lesser extent also the midlegs, the stepping frequency even increased upon amputation, as if they were continually seeking surface contact (Fig. 1j).

We conclude from this series of experiments that MDN induces backward walking predominantly by engaging the muscles that control the hindleg coxa-trochanter and femur-tibia joints. MDN switches the phases of flexion and extension for these joints during the stepping cycle, and enhances their movement in the hindleg so as to mirror those of the corresponding foreleg joints during forward walking. As a result of these adjustments to hindleg stepping, it is the hindlegs that provide the main propulsive force during backward walking. MDN also provides coordinated neural input to the midleg and foreleg motor circuits. However, MDN’s impact in these segments is considerably weaker and, in the absence of any drive from the hindlegs, insufficient to propel the animal backwards.

### Trans-Tango reveals cells postsynaptic to MDNs

Having determined how MDN activation changes leg kinematics, we next sought to trace the neural circuits through which MDN effects these changes. In a first approach, we used the trans-Tango method^16^ to identify cells in the VNC likely to be post-synaptic to MDNs. Trans-Tango employs a heterologous ligand to activate its cognate receptor in postsynaptic cells, which then become labelled by expression of a QF-dependent reporter. We targeted the trans-Tango ligand to MDNs and the trans-Tango receptor to all neurons, and acquired confocal images of the VNC to visualize post-synaptic cells labelled by an mtdTomato-3HA reporter (Fig. 2a). A large number of cells in the VNC were MDN trans-Tango-positive, and, as expected, their neurites overlap extensively with the axonal arborizations of the MDNs in all three leg neuropils (Fig. 2a). This high density of trans-Tango labelling precluded the identification of individual cell types. In order to visualize single trans-Tango-positive cells, we therefore used a stochastic variant of trans-Tango ^16^ in which only a small subset of the postsynaptic cells were labelled in any one sample (Fig. 2b). A total of 541 samples were imaged by confocal microscopy and registered onto a common reference template (ref. 17; Fig. 2b).

**Fig. 2.**
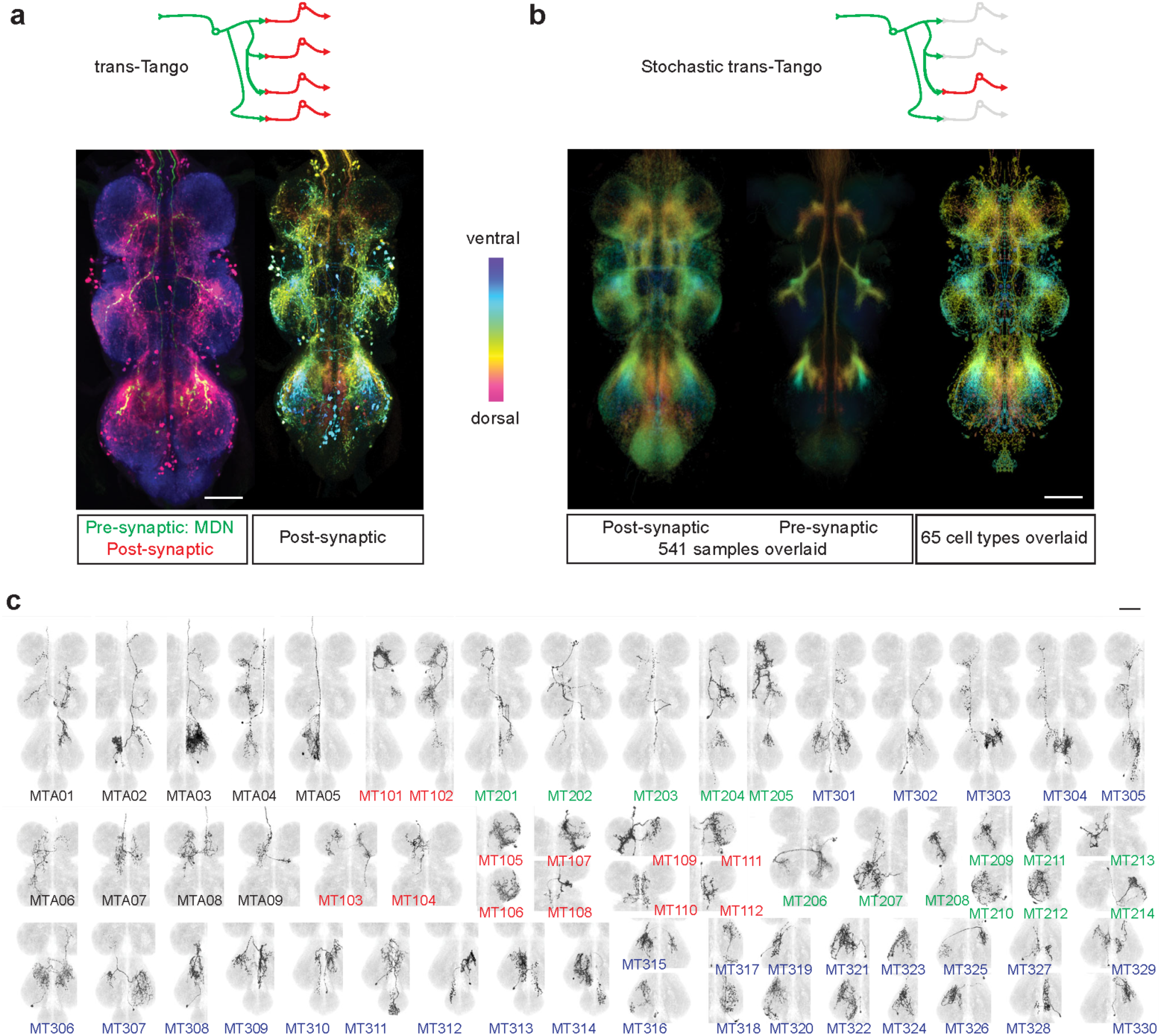
Trans-Tango reveals cells postsynaptic to MDNs. a, Top, schematic of trans-Tango labelling, showing target cell in green and trans-synaptically labelled cells in red. Bottom, an example of trans-Tango labelling of cells postsynaptic to MDNs, targeted with *MDN-1-GAL4*. Left, maximal projection confocal image of the VNC, showing MDN axons in green (anti-GFP), trans-Tango labelled cells in red (anti-HA), and all synapses in blue (anti-Bruchpilot, nc82). Right, the same sample registered onto a standard VNC template, with the trans-Tango channel shown as a color-coded maximal intensity projection (colorMIP) in which color encodes the z-section in which the maximum occurs. b, Top, schematic of stochastic trans-Tango labelling. Bottom left and middle, standard deviation projections of colorMIP images of the post- and pre-synaptic channels, respectively, of 541 stochastic trans-Tango samples. Bottom right, standard deviation projection of 65 types of segmented neurons and their mirror images. c, Segmented MT cell types from stochastic trans-Tango dataset, shown as registered maximum intensity projections overlaid on the JRC2017 template VNC. Cell type designations are colored to indicate ascending neurons (black), and interneurons with soma located in either T1 (red), T2 (green), or T3 (blue). Scale bars: 50 μm.

The cell types labelled by trans-Tango were highly diverse in their morphology. We identified 65 morphologically distinct cell types amongst the 541 samples, and segmented a representative cell of each class (Fig. 2c). Collectively, these neurons appear to encompass the full pattern of trans-Tango-positive cells (Fig. 2b). These cell types comprise 9 classes of ascending neurons and 56 interneurons. We did not detect any motor neurons directly post-synaptic to the MDNs. The majority of these putative MDN output cells (4/9 ascending neurons and 30/56 interneurons) have their soma located in the T3 segment, consistent with the inference from the joint kinematic analysis that MDN predominantly impacts the T3 motor circuits. For the purposes of this work, we assigned these cell types provisional names starting with MT (for “MDN trans-Tango”), followed by either an A for ascending neurons or 1, 2, or 3 for interneurons in the T1, T2 or T3 segments, respectively, and finally a consecutive two-digit number (Fig. 2c).

### Volumetric calcium imaging reveals cell types responsive to MDN activation

Labeling with trans-Tango provides no information as to the strength or sign of synaptic connections, nor can it reveal cell types that are functionally responsive to the MDNs but not amongst their immediate post-synaptic partners. We therefore complemented this anatomical approach with volumetric functional imaging (Fig. 3a), optogenetically activating the MDNs using CsChrimson^16^ and monitoring calcium responses at 1 Hz in the VNC with GCaMP6s^19^. MDN was activated with a series of 20-s red light pulses, one per minute over a 10-minute period. Responsive cells were identified using a voxel-wise analysis of covariance with a kernel that captures the dynamics of the stimulus protocol and GCaMP6s response kinetics (Fig. 3a). In initial experiments in which GCaMP6s was expressed in all neurons, or all glutamatergic neurons, we reproducibly observed both excitatory and inhibitory responses in the VNC, predominantly but not exclusively in the T3 segment (Fig. 3b).

**Fig. 3.**
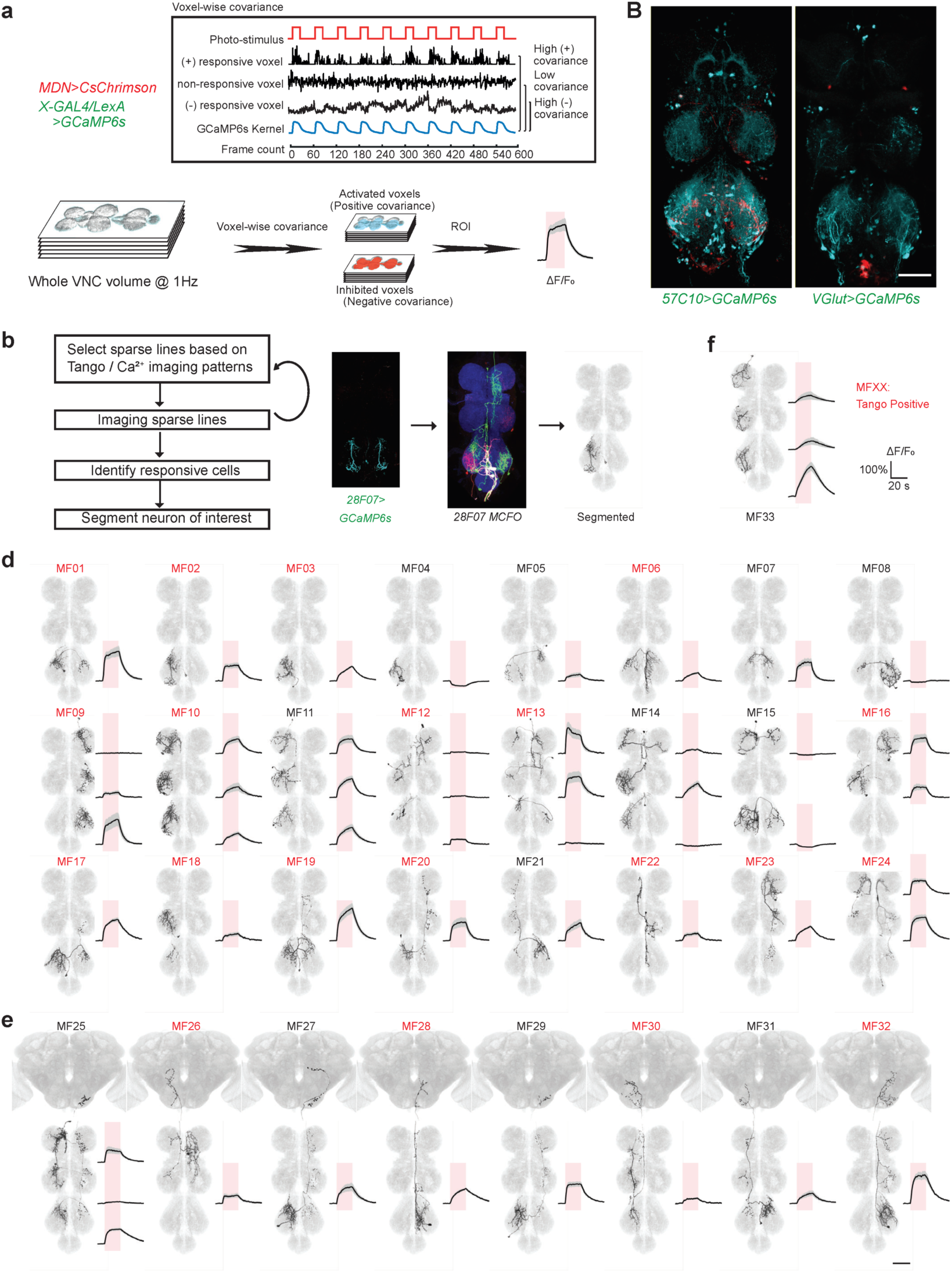
MDN-responsive neurons identified by functional imaging. a, Schematic of the functional imaging and data analysis pipeline. b, Maximum intensity projections of imaged VNC volumes during photostimulation, using *MDN-1-GAL4* to drive Chrimson88 in MDNs. Cyan represents activated voxels; red represents inhibited voxels. Scale bar: 50 μm c, Left, pipeline for discovering cell types that are functionally connected to MDN. Right, an example showing how an MDN-responsive cell type is identified. The three images are, from left to right, a maximum intensity projection of activated voxels in the VNC of a fly in which GCaMP6s expression is driven by *28F07-LexA*, a multi-color flip-out (MFCO) image from the corresponding *28F07-GAL4* driver, and a single cell segmented from the MCFO image that matches the morphology of the MDN-responsive neurons. d, Twenty-four MF interneuron types, whereby segmentally repeated cells are classified as the same cell type. Maximum intensity projections of segmented and registered images are shown on the left; averaged responses to MDN activation on the right (mean ± s.e.m. of *n* ≥ 5 samples; red shading indicates the 20 s red light stimulus; see also Extended Data Fig. 2). Cell types labelled in red were also identified in the trans-Tango experiments (see Extended Data Fig. 1 for pairwise comparisons). e, Eight MF ascending neuron types and their calcium responses to MDN activation. f, The single MF motor neuron type and its calcium response to MDN activation.

In functional imaging experiments in which GCaMP6s was broadly expressed, it was not possible to reliably identify individual responsive cells. We therefore used an iterative approach in which we screened selected lines from large collections of GAL4 or LexA drivers^20,21^. We started with a panel of relatively broadly expressed lines, and then, for any line in which we detected MDN-responsive cells, iteratively examined successively sparser lines likely to contain these cells. Additionally, we included sparse driver lines likely to include any of the cell types identified in the trans-Tango experiments. For the sparsest lines in which a GCaMP6s response was observed, we identified the specific cell type that was MDN-responsive by aligning the pattern of correlated voxels to a series of images of single cells obtained by stochastic labelling22 using a driver line containing the same enhancer (Fig. 3c). These identified neurons were then manually segmented from the stochastically labelled samples.

While this iterative sampling approach inevitably fails to isolate all the MDN-responsive cells, it was sufficiently broad in scope to allow us to resolve a total of 33 morphologically distinct MDN-responsive cell types (Fig. 3d-f). We provisionally refer to these cell types here as MF cells (MDN functional imaging). As for the MT cells identified with trans-Tango, the MF cells also include both interneurons (Fig. 3d) and ascending neurons (Fig. 3e), and are predominantly located in T3. Additionally, one type of MDN-responsive motor neuron was identified in the functional imaging experiments (Fig. 3f). Not surprisingly, and indeed in part due to our search strategy, there is considerable overlap amongst the cells identified by trans-Tango and by functional imaging, with a total of 20 cell types detected by both methods (Extended Data Fig. 1).

The MF cells exhibit a diverse array of responses to MDN activation (Fig. 3d-f and Extended Data Fig. 2). Most MF cell types showed a positive calcium response upon MDN activation, as expected given that the MDNs are cholinergic, and hence excitatory, and GCaMP6s more reliably reports excitation than inhibition. Some cells however consistently showed inhibitory responses to MDN activation. None of these inhibited cells were trans-Tango positive, and indeed would not be expected to be directly post-synaptic to the MDNs. Both the excitatory and inhibitory responses varied considerably in their amplitude and dynamics, even amongst segmentally repeated cells of the same type. For example, the MF09, MF13, and MF25 cells are each present in all three segments, yet MDN activation elicited strong activation only in the T3 MF09s, the T1 and T2 MF13s, and the T1 and T3 MF25 (Fig. 3d). An even more dramatic example of segment-specific MDN responses is MF14, for which the T1 and T2 cells were excited but the T3 cells inhibited (Fig. 3d).

### Several MF cell types contribute to MDN-induced backward walking

Even the sparsest driver lines we used to identify the MF cells in the functional imaging experiments typically label several additional cell types. Although these other cell types are not MDN-responsive, they may nonetheless perform critical functions during walking. Thus, this initial set of GAL4 driver lines was not ideal for a functional analysis of the MF cells in backward walking. We therefore used the split-GAL4 method^21,23,24^ to generate highly-specific driver lines for selected MF cells. These lines were generated as part of a larger effort to systematically classify and genetically target each of the neuronal cell types in the leg neuropils (R.M., K.F., B.J.D., in preparation). We matched each of the segmented MF cell profiles to specific cell types defined in this systematic classification, thereby identifying a total of 82 stable split-GAL4 (*SS*) lines that collectively covered 29 of the 33 MF cell types (Fig. 4a and Extended Data Fig. 3a).

**Fig. 4.**
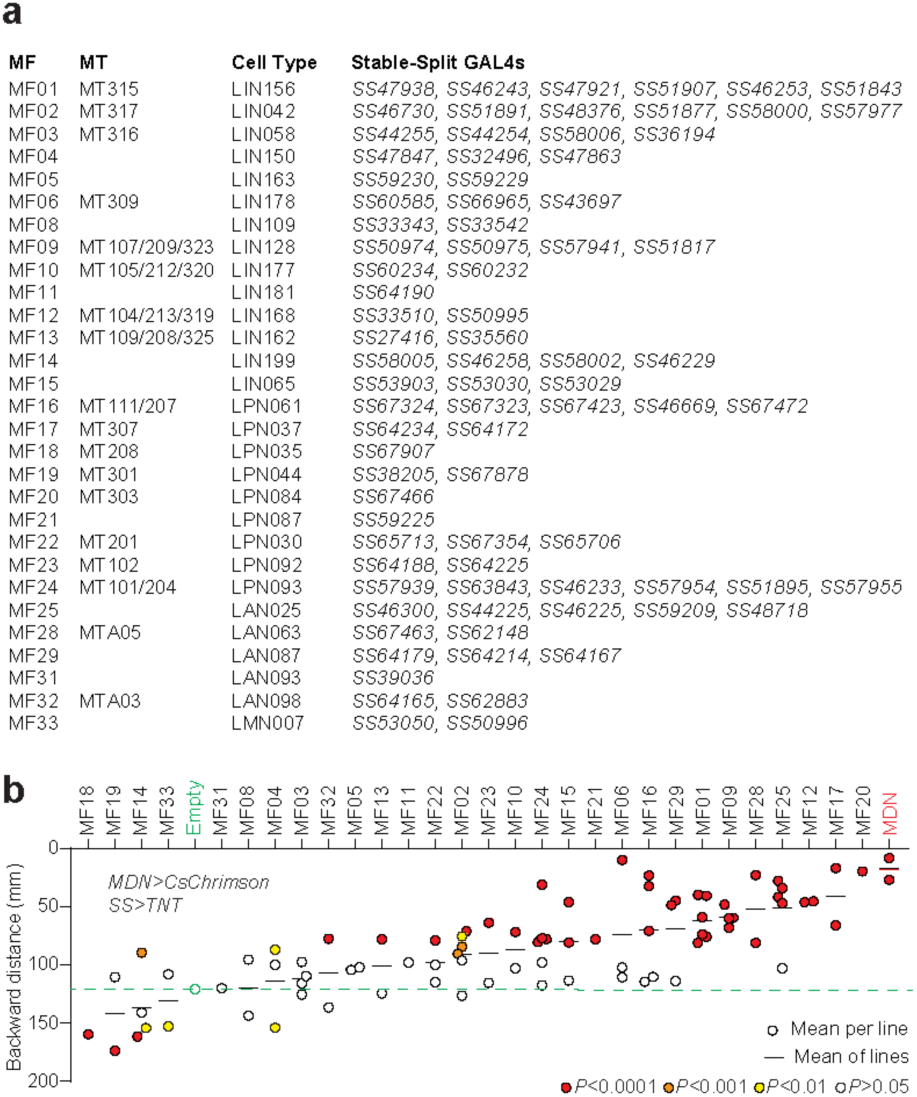
Several MF cell types contribute to MDN-induced backward walking. a, Correspondence between 29 MF cell types, their matching MT cell types, cell types identified in a systematic analysis of leg neuropil neurons, and the split-GAL4 lines that label them and were used to generate the data shown in (b). b, Total backward walking distance during 9 episodes of a 5-s photoactivation of MDN, in flies in which a single MF cell type was silenced. Each data point represents the mean of all flies tested with a single split-GAL4 line, shaded to indicate *P* values from one-way ANOVA tests with post-hoc Dunnett’s correction for multiple comparisons. For cell types for which multiple split-GAL4 lines were tested, the mean across all lines is shown. Dashed green dashed line marks the backward walking distance of negative control flies, for comparison. See also Extended Data Fig. 3b.

We then used these split-GAL4 driver lines to determine whether genetically silencing each class of MF cell was able to suppress MDN-induced backward walking in an open arena (FlyBowl^25^). For this we generated flies in which CsChrimson was expressed in MDNs using a split-LexA driver, and tetanus toxin light chain^26^ in a single class of MF cells using one of the *SS* lines (Extended Data Fig. 4). As a negative control we used an empty GAL4 driver (lacking a neuronal enhancer element); as positive controls we used two different MDN GAL4 drivers^10^. For each of these genotypes, we typically examined a total of approximately 50 flies distributed across 2 trials for their locomotion responses to red light stimulation in an open arena (Extended Data Fig. 3b). Fig. 4b shows the mean backward locomotion during the stimulus period for each of the 82 MF split-GAL4 lines and the 3 controls. For several MF cell types, multiple driver lines all resulted in a strong impairment of backward walking. Some of these also exhibited defects in spontaneous forward walking (Extended Data Fig. 3b), but many specifically disrupted backward walking. Of these, we decided to focus our further analysis on two of the cell types for which we had the most *SS* driver lines and the most consistent reduction of backward walking: MF01, which corresponds to the LIN156 cells in our systematic classification of leg neuropil neurons, and MF09, which corresponds to LIN128. For these two cells types, a total of 6 and 4 *SS* driver lines, respectively, were all able to suppress MDN-induced backward walking. We next asked if and how each of these two cell types contributes to MDN’s effect on hindleg joint movements.

### LIN156 neurons trigger tibia flexion via the tibia-reductor motor neurons

The LIN156 cells, identified in both the MDN trans-Tango (MT315, Fig. 2c) and functional imaging experiments (MF01, Fig. 3d), are a bilateral pair of neurons specific to the T3 neuropils (Fig. 5a). They resemble the only surviving neurons of the 14B lineage^27^. Each of 6 LIN156 split-GAL4 driver lines reduced MDN-induced backward walking in the neuronal silencing experiments (Fig. 4b). The most restricted of these lines are *SS47938* and *SS46243. SS47938* labels no other cells in the central nervous system, whereas *SS46243* additionally labels a pair of ascending neurons (Extended Data Fig. 3a). Using these two cell-type-specific driver lines in calcium imaging experiments, we confirmed that the LIN156 neurons indeed respond to MDN activation (Fig. 5b).

**Fig. 5.**
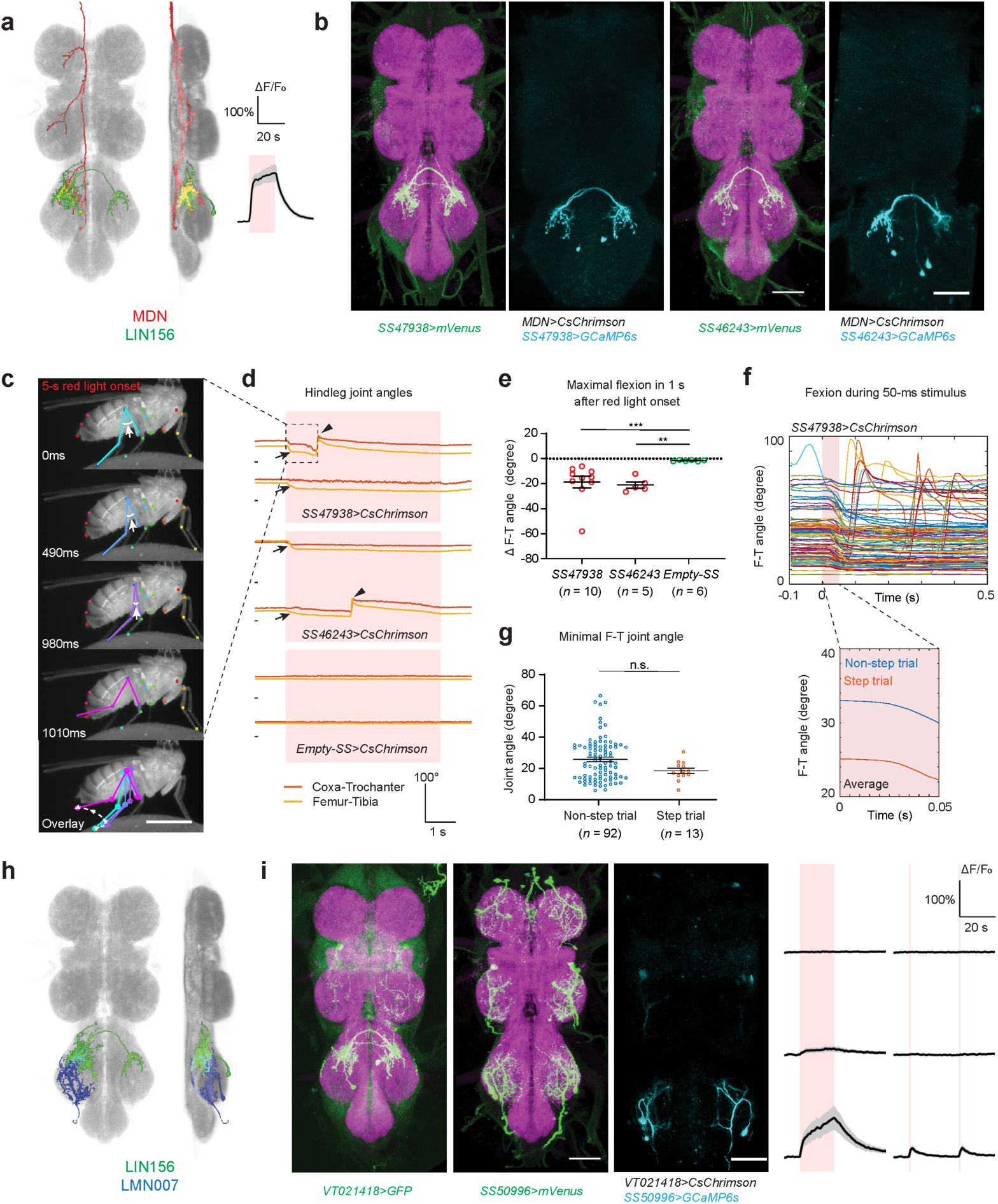
LIN156 neurons control tibia flexion. a, Segmented and registered images of MDN (red) and LIN156 (green) from ventral and lateral perspectives, and calcium response in LIN156 upon photoactivation of MDN. Same data as in Fig. 3d. b, Confocal images of VNCs of two split-GAL4 lines labelling LIN156 (magenta, nc82 staining; green, anti-GFP staining of CsChrimson-mVenus reporter), and activated voxels (cyan) from calcium imaging experiments using each line. Scale bars: 50 μm. c, Selected frames from a representative video of LIN156 activation in a tethered, decapitated fly, during a 5-s red light stimulus from t=0 ms. Arrows indicate the femur-tibia angles. Dashed line with arrows in the overlay image indicates sequence of movement. Scale bar: 1 mm. See also Supplementary Video 3. d, Hindleg joint angle time series, showing two representative traces for each genotype. Red shade indicates a 5-s pulse of red light. A short black line marks the origin for each trace (t=0 s, 0 degrees). Arrows indicate the initial femur-tibia flexion; arrowheads mark steps. The top traces are from the video shown in (c). e, Maximal femur-tibia joint flexion within 1 s after red light onset. ***, *P*<0.001; **, *P*<0.01, Mann-Whitney test, two-tailed *P* values. f, Top, overlaid hindleg femur-tibia joint angles from trials in which LIN156 was activated by a 50 ms light pulse. Bottom, mean joint angle traces, calculated separately for trials with and without a step. Red shading indicates the red light stimulus. *N* = 3 flies and *n* = 105 trials. g, Minimum femur-tibia joint angles for non-step and step trials. n.s., *P*≥0.05, Mann-Whitney test. h, Registered segmented image of LIN156 (green), LMN007 (blue), and all synapses (gray, nc82). i, Confocal images of VNCs of a *LexA* line labelling LIN156 and *SS50996*, labelling LMN007 (green, anti-GFP staining of a myr-GFP or CsChrimson-mVenus reporter; magenta, nc82), and activated voxels (cyan) from calcium imaging of LMN007 upon LIN156 activation. Traces show the averaged responses for LMN007 to LIN156 activation (mean ± s.e.m., *N* = 6 flies), upon either a single 20-s stimulus or two 1-s light pulses (red shading).

If the LIN156 neurons mediate specific joint movements during MDN-induced backward walking, then optogenetic activation of these neurons in stationary flies might elicit these joint movements in isolation. We tested this prediction using tethered, decapitated flies. Decapitation removes all descending inputs to the leg motor circuits, so that any leg movements induced by optogenetic activation of leg interneurons are not obscured by movements triggered by brain activity. A tethered, decapitated fly is normally stationary, but optogenetic activation of the severed VNC projections of the MDNs induces backward walking, with its characteristically exaggerated hindleg movements (Extended Data Fig. 5).

In this decapitated fly preparation, optogenetic activation of LIN156 using either *SS47938* or *SS46243* reliably induced hindleg tibia flexion at stimulus onset (Figs 5c, d [arrows], and e). As the stimulus persisted, the initial tibia flexion was occasionally followed by rapid leg lifting and extension to execute a full backward step cycle (arrowheads in Fig. 5d). A similar pattern of tibia flexion, followed in some cases by stepping, was also seen in a series of experiments using a shorter 50 ms stimulus (Fig. 5f and Supplementary Video 3). With this shorter stimulus, the stepping occurred with a variable delay and outside the stimulus window, suggesting that it is not a direct consequence of LIN156 activation. If stepping occurred, it was initiated from a small joint angle (Fig. 5g). However, there was no significant difference in the minimum angle reached in stepping versus non-stepping trials, suggesting that reaching this small angle does not automatically trigger the next swing phase (Fig. 5g).

These photoactivation experiments suggest that LIN156 is a premotor neuron for tibia flexion. The single class of motorneuron identified in our functional imaging experiments, MF33, has a morphology that matches that of the motor neurons innervating the tibia reductor muscle^28,29^, which is responsible for tibia flexion^30,31^. In our systematic classification of neurons in the leg neuropils, we designated these as the LMN007 neurons (R.M., K.F., and B.J.D., in preparation). They are present in each segment, but upon MDN stimulation are preferentially activated in T3 (Fig. 3f). We confirmed that the LMN007 neurons are indeed the tibia reductor motor neurons by examining their muscle innervation in the leg (Extended Data Fig. 6a). As expected, optogenetic activation of the LMZ007 neurons induced hindleg tibia flexion, albeit without the re-extension that was observed with LIN156 activation (Extended Data Fig. 6b-d). Calcium imaging experiments demonstrated that the T3 LMN007 neurons are activated upon stimulation of either the LIN156 neurons (Fig. 5h,i) or the MDNs (Extended Data Fig. 6e), and a GRASP experiment^32^ suggested that they are direct synaptic partners of the LIN156 cells (Extended Data Fig. 6f).

We conclude from these experiments that LIN156 activates the LMN007 neurons to induce hindleg tibia flexion. LIN156 and LMN007 are thus candidates to provide the hindleg power stroke during the stance phase of backward walking.

### LIN128 neurons trigger leg lifting

Like the LIN156 neurons, the LIN128 neurons were also identified in both the MDN trans-Tango (MT323, Fig. 2c) and functional imaging experiments (MF09, Fig. 3d), and are also required for robust MDN-induced backward walking (Fig. 4b). LIN128 neurons are segmentally repeated local interneurons, but were most strongly activated by MDN in T3 (Fig. 6a). The T2 LIN128 neurons responded less strongly to MDN activation, and no response was detected in the T1 LIN128s. The most restricted split-GAL4 lines we obtained for these neurons are *SS50974* and *SS50975. SS50974* labels only the T3 cells, whereas *SS50975* labels cells in all three thoracic segments (Fig. 6b). Neither driver labels any other cells in the CNS (Extended Data Fig. 3a). We confirmed with these drivers that both the T2 and T3 LIN128 neurons are activated upon optogenetic stimulation of the MDNs (Fig. 6b; note that *SS50975* drives weaker expression in T3 than in T1 and T2).

**Fig. 6.**
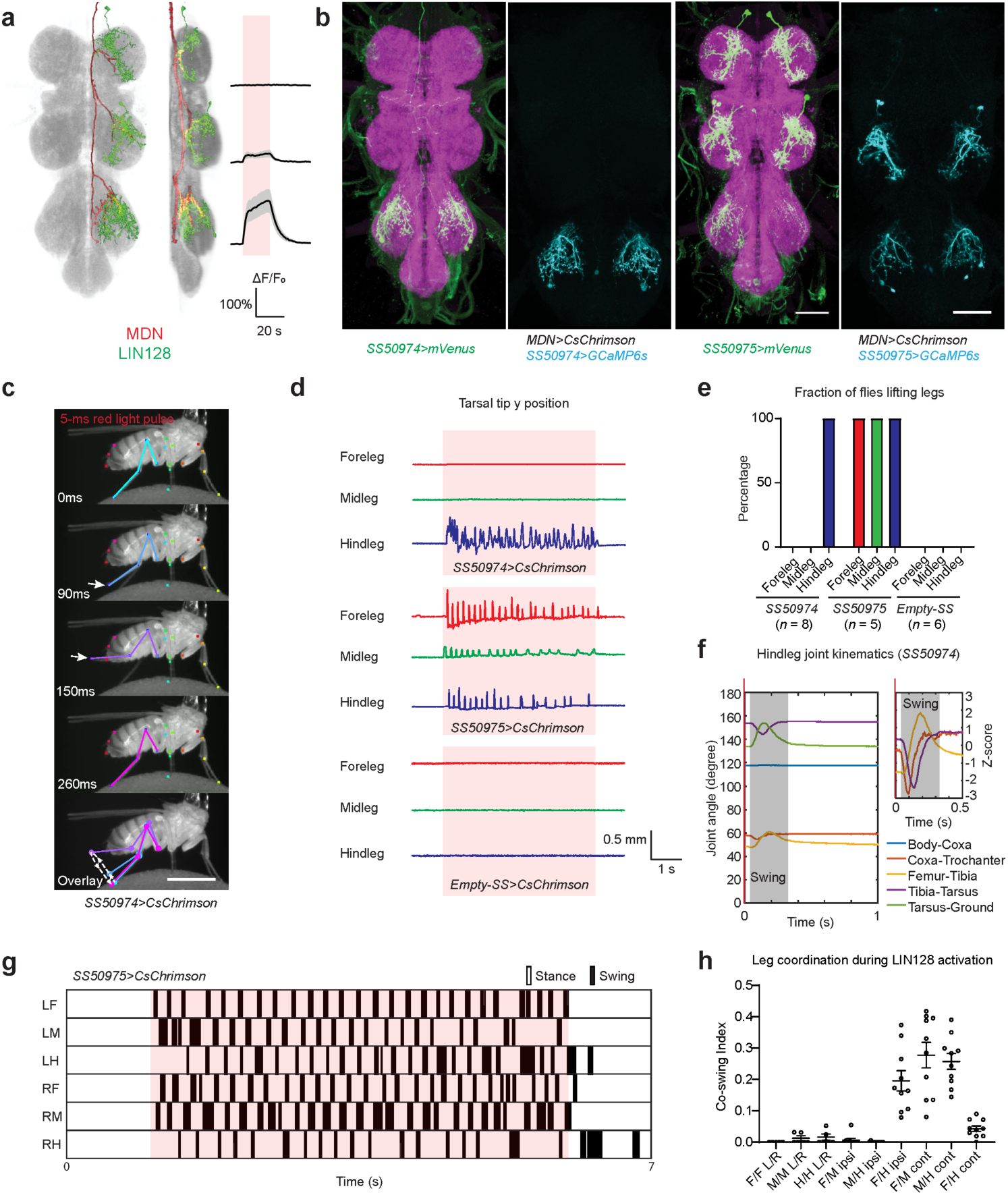
LIN128 neurons trigger leg lifting. a, Segmented and registered images of MDN (red) and LIN128 (green) from ventral and lateral perspectives, and calcium responses of LIN128 upon MDN activation. Same data as in Fig. 3d. b, Confocal images of VNCs of two split-GAL4 lines labelling LIN128 (magenta, nc82 staining; green, anti-GFP staining of CsChrimson-mVenus reporter), and activated voxels (cyan) from calcium imaging experiments using each line. Scale bars: 50 μm. c, Selected frames from a representative video of LIN128 activation in a tethered, decapitated fly upon a 50-ms red light stimulus applied at t = 0 ms. Arrows indicate the femur-tibia angles. Dashed line with arrows in the overlay image indicates sequence of movement. Scale bar: 1 mm. See also Supplementary Video 4. d, Representative traces of tarsal tip positions for each leg. Red shade indicates a 5-s pulse of red light. e, Percentage of flies lifting each leg. f, Left, averaged hindleg joint angles upon photoactivation of LIN128 by 5-ms red-light pulses, starting at 0 ms. Grey indicates the inferred swing phase. Right, z-scores of coxa-trochanter, femur-tibia and tibia-tarsus angles from the same data, highlighting the phase relationships between joints. *N* = 3 flies and *n* = 100 trials. g, Representative footfall pattern during a continuous 5-s photoactivation of LIN128 (red shading). LF, LM, LR, RF, RM, RH designate left and right fore-, mid- and hindlegs, respectively. See also Supplementary Video 5. h, Leg coordination during 5-s photoactivation of LIN128, quantified as a co-swing index (the total time both legs are in swing divided by the time either leg is in swing). *N* = 5 flies (*n* = 5 intrasegmental leg pairs, *n* = 10 intersegmental pairs). F, foreleg; M, midleg; H, hindleg; L, left; R; right; ipsi, ipsilateral; cont, contralateral.

Using decapitated and tethered flies, we found that optogenetic activation of LIN128 induces lifting and a swing-like movement of the leg. With the *SS50974* driver, only the hindlegs were lifted (Fig. 6c-e and Supplementary Video 4); with *SS50975*, all 6 legs were lifted (Fig. 6d and e). We examined the hindleg lifting induced by *SS50974* activation more closely and observed that a single brief (5 ms) pulse of red light stimulation elicited a coordinated sequence of leg movements involving all three joints and resembling the swing phase of backward walking (Fig. 6f). Using the *SS50975* driver to activate the LIN128 neurons in all three segments, we found that a longer red light stimulus (5s) resulted in an alternating pattern of stepping across all 6 legs. Despite the simultaneous optogenetic activation of all LIN128 neurons, the legs stepped in a coordinated pattern similar to that typically observed in forward walking, with the fore- and hinglegs on one side of the body in phase with the midlegs on the other (Fig. 6g,h and Supplementary Video 5; refs.33,34). These data suggest that the LIN128 neurons initiate swing phase, which then follows the normal pattern of both inter-joint and inter-leg coordination.

### LIN156 and LIN128 function in the stance and swing phases of backward walking

Finally, we sought to test the functional hypotheses we derived from the activation experiments: that LIN156 and LIN128 act at distinct timepoints during the backward stepping cycle, with LIN156 providing the hindleg power stroke during stance phase and LIN128 neurons subsequently initiating swing phase. We tested these hypotheses by silencing either the LIN156 or LIN128 cells in tethered flies induced to walk backwards by MDN activation, and tracking the resulting hindleg joint movements. Such experiments required an optogenetic silencer and an activator that could be independently controlled with minimal crosstalk. We found that we could best achieve this by stimulating CsChrimson with 660nm red light and the neuronal silencer GtACR2 (ref. 35) with 470nm blue light. With these tools, we established a protocol using tethered and decapitated flies in which CsChrimson was expressed in the MDNs and GtACR2 in either the LIN156 or LIN128 neurons. Positive control flies expressed GtACR2 in the MDNs; negative controls used an “empty” split-*GAL4* driver that resulted in no GtACR2 expression. Each fly was subject to a series of control and experimental trials (Fig. 7a). Control trials consisted solely of a 25-s red-light stimulus to activate the MDNs. Experimental trials additionally included a 10-s blue light stimulus to silence the LIN156 or LIN128 neurons, applied 10-s after the onset of the red-light stimulus.

**Fig. 7.**
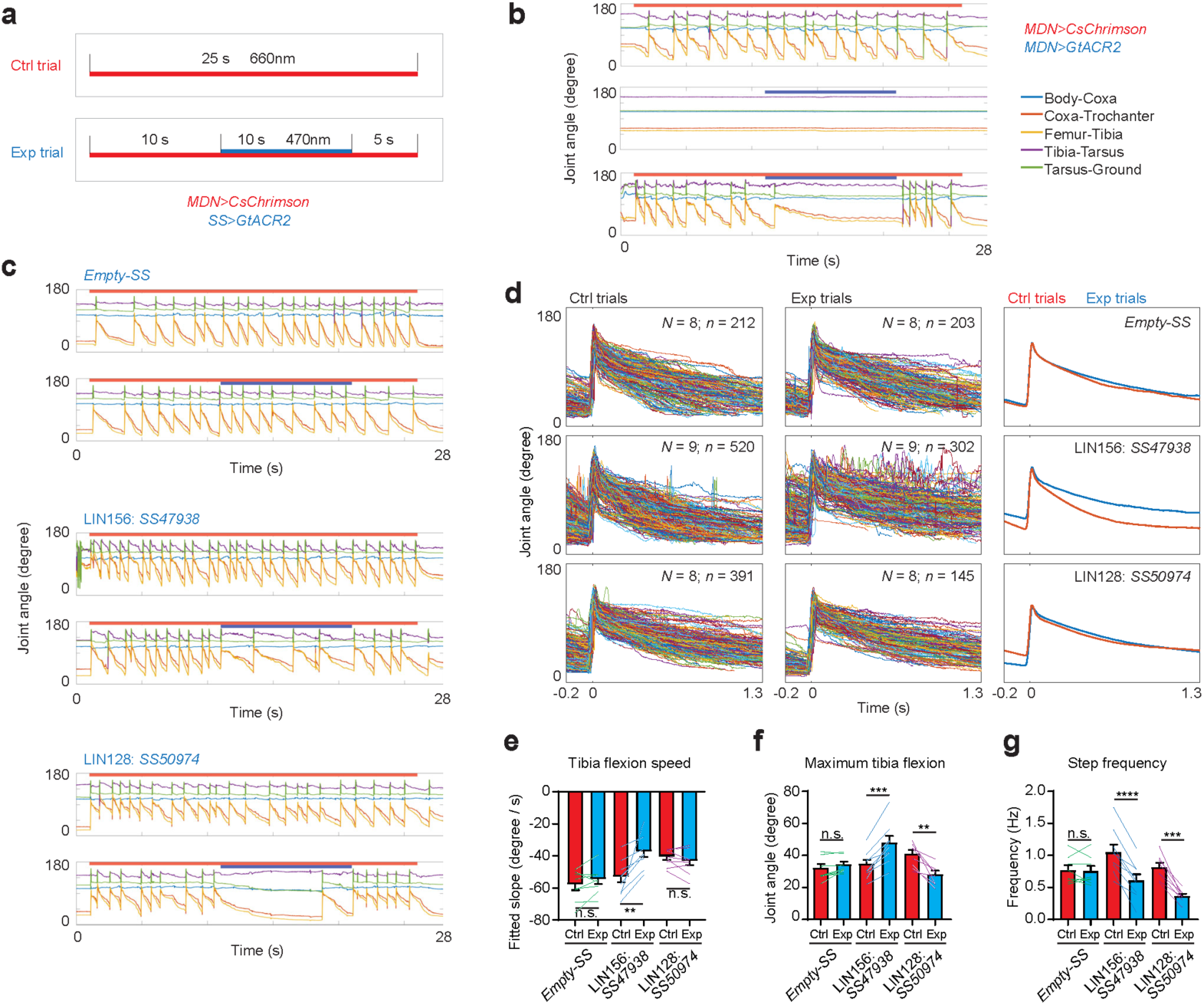
LIN156 and LIN128 function in the stance and swing phases of backward walking. a, Experimental protocol. Each fly was subject to the same number of control (Ctrl) and experimental (Exp) trials. Red bar indicates red-light stimulus; blue bar indicates blue-light stimulus. b, Representative hindleg joint angle time series for positive control flies expressing both CsChrimson and GtACR2 in MDN, and stimulated with either red and blue light alone, or both. c, Representative hindleg joint angle time series for flies of the indicated genotypes. See also Supplementary Videos 6 and 7. d, Left and middle: overlaid femur-tibia joint angle from all steps during the 10-s blue-light stimulus in experimental trials, and the corresponding period in control flies. Right: average of all steps for experimental (blue) and control trials (red). Before averaging, individual traces were truncated if another step occurred within the 1.5 s time window. e-g, Slope of femur-tibia joint angle change (e) and minimum angle (f) during stance phase, and step frequencies (g) during the 10-s blue light stimulus period of experimental trials and the corresponding period of control trials. All data are shown as mean ± s.e.m. across all *N* flies, with colored lines pairing experimental and control values for the same fly. ****, *P*<0.0001; ***, *P*<0.001; **, *P*<0.01; n.s., *P*≥0.05; paired t-tests, two-tailed *P* values.

In positive control flies, MDN-induced backward walking was fully suppressed by optogenetic silencing of the MDNs themselves (Fig. 7b). Conversely, negative control flies were unaffected by the blue light stimulus in experimental trials (Fig. 7c). When LIN156 neurons were silenced, the power stroke appeared to be weakened, as suggested by the reduced speed and amplitude of tibia flexion, and hence a slight delay in the initiation of swing (Fig. 7c and Supplementary Video 6). In contrast, when LIN128 neurons were silenced, the power stroke was unaffected, as judged by the initial speed of tibia flexion. However, in these LIN128-silenced flies, the leg appeared to be stuck in stance phase, with the tibia flexing to a much greater extent before swing was eventually initiated (Fig. 7c and Supplementary Video 7).

We quantified these effects by measuring the femur-tibia joint angle during each step in the 10-s blue-light stimulus period of experimental trials and the corresponding periods of control trials for the same fly (Fig. 7d). We aligned these curves by time-point of their maxima, which typically occurred in the middle of the short swing phase, 20–30 ms before the start of stance phase. From these curves we then determined, for each fly, the average initial speed and maximum extent of tibia flexion during stance phase, and the frequency of stepping, across all experimental and control trials (Fig. 7e-g). This analysis confirmed our initial impression that LIN156-silenced flies, but not LIN128-silenced flies, had a reduced speed of tibia flexion during the blue-light stimulus (Fig. 7e). In LIN156-silenced flies, the femur-tibia joint was less flexed than in negative control flies; in LIN128-silenced flies it was more flexed (Fig. 7f). Both LIN156- and LIN128-silenced flies took fewer steps (Fig. 7g), presumably however for different reasons. In LIN156-silenced flies, the late transition to swing phase is explained by a slower stroke (Fig. 7e), not a greater degree of flexion (Fig. 7f). In LIN128-silenced flies, the late initiation of swing correlated with a greater flexion (Fig. 7f), not a slower stroke (Fig. 7e). Similar results were obtained when we compared the blue-light stimulus period of experimental trials to the pre- or post-blue-light stimulus periods of the same trials, rather than to control trials (Extended Data Fig. 7a and b). We also found that the increase in hindleg stepping that results from tarsal amputation (Fig. 1i and j) is similarly suppressed by LIN128 silencing, suggesting that these oscillations, like normal steps, are centrally driven (Extended Data Fig. 7c).

These data thus confirm our hypotheses from the activation experiments. LIN156 and LIN128 neurons mediate two distinct aspects of MDN-induced backward walking. LIN156 neurons function in tibia flexion, providing the power stroke during stance phase. LIN128 neurons facilitate leg lifting at the end of stance phase to initiate swing and complete the stepping cycle.

## Discussion

Backward walking triggered by the MDNs is an excellent model for exploring how descending command-like neurons produce abrupt but coordinated changes in motor circuits. The data presented here indicate that MDN acts primarily on the T3 motor circuits, such that the hindlegs become the major driving force in backward walking. The T1 and T2 motor circuits also receive MDN input, but the backward stepping of the fore- and midlegs is primarily dependent upon proprioceptive signals generated as the hindlegs propel the body backwards. Through a combination of trans-synaptic tracing and functional imaging, we identified over 30 neuronal cell types targeted by MDN – the majority of them in T3. Amongst these are the LIN156 and LIN128 neurons, which act at separate stages in the hindleg stepping cycle. LIN156, a premotor neuron for tibia flexion, mediates the power stroke in stance phase. LIN128, an interneuron that triggers leg lifting, is responsible for the initiation of swing. Together, these data suggest that the MDNs do not act solely, if at all, through a high-level bistable switch that controls the stepping direction ^36,37^, but rather individually modulate each of the diverse motor outputs needed for coordinated backward walking.

### Stance phase

Tibia flexion and femur elevation are the most prominent hindleg joint movements during the stance phase of backward walking – the two movements largely occurring in synchrony to fold up the leg, thereby pulling the body backwards. The LIN156 neurons, which are specific to T3, flex the hindleg tibia by activating LMN007, one of the tibia flexor motor neurons. Femur elevation may be mediated through other MDN-effector neurons, possibly included amongst the MF and MT cell types we have not yet examined in detail. There may also be some mechanical coupling between the two joints, as we often observed femur elevation (flexion of the coxa-trochanter joint) when tibia flexion was induced by activating either LIN156 or LMN007. Such mechanical coupling between tibia flexion and femur elevation has been observed in the stick insect^38^. Femur elevation was however generally weaker and less reliable than tibia flexion in the LIN156 activation experiments, suggesting that it is not the primary function of these neurons.

LMN007 is one of a large group of motor neurons that innervate the tibia flexor muscles. It most closely resembles the “slow” tibia flexor motor neurons, which have the lowest activation threshold and highest spiking rate but generate the weakest force31. It is possible that LIN156 also activates other tibia flexor motor neurons, but the *ex vivo* nature of our preparation or the limited sensitivity of our imaging method hampered our ability to detect these responses. Indeed, silencing of LIN156 suppressed MDN-induced backward walking, whereas silencing of LMN007 did not. This suggests that at least some other tibia flexor motor neurons may also be involved in backward walking.

In behavioral assays, the fly typically backs up for several steps when it encounters an obstacle, despite receiving only a transient mechanosensory stimulus. However, upon artificial activation of MDNs, the fly walks backwards only so long as the stimulus persists^10,11^. These results suggest that circuits in the brain convert an acute mechanosensory stimulus into tonic activation of MDNs. This conclusion is supported by *in vivo* functional imaging experiments, which showed a sustained calcium response in the MDNs during bouts of backward walking^39^. The LIN156 and LMN007 neurons, in contrast, are presumably only phasically active, becoming silent when the leg pauses or is re-extended. The timing of LIN156 and LMN007 activation must therefore be determined by other inputs, most likely including proprioceptive signals from the same as well as other legs^40-42^. One such input may be phasic inhibition from proprioceptive neurons that are activated when the tibia nears full flexion ^43^.

### Swing phase

To complete the stepping cycle, the retracted leg must initiate swing and return to its extended position. Several factors act together to initiate swing, including the position of the leg, the unloading of the leg, and the loading of other legs^40,41,44-46^. Some or all of these factors might lead to activation of the LIN128 neurons, which trigger leg lifting and initiate swing phase. It is interesting to note that the LIN128 neurons are present in all three segments but are most strongly activated by MDN in T3. Perhaps, during backward walking, the relative weighting or timing of the factors that initiate swing are most dramatically altered for the hindleg, and MDN effects these changes. For example, when walking forwards, the hindleg initiates swing from an extended position; when walking backwards, swing is initiated from the folded position. Further insight into the circuit mechanisms that initiate swing should come from mapping proprioceptive and other inputs to LIN128 neurons, and determining how MDN activity shapes LIN128’s responses to these inputs. This circuit mapping may also help elucidate the role of LIN128 in interleg coordination. The recent release^47^ of an electron-microscope volume of the VNC should greatly facilitate these circuit reconstruction efforts.

We propose that, as for LIN156, it is primarily the intra- and interleg sensory feedback signals that ensure the phasic activation of LIN128 neurons despite tonic input from MDNs. But it is also possible that inhibitory pathways entirely within the central motor circuits also contribute to the alternating phasic activation of LIN156 and LIN128. These central mechanisms might include reciprocal feedforward inhibition. Feedforward inhibition from LIN128 to LIN156 could relax the tibia flexor during the return stroke, while inhibition from LIN156 to LIN128 may help keep leg grounded throughout its power stroke. In this scenario, LIN156 and LIN128 may be intergral components of a central pattern generator for stepping, the neuronal basis of which has been elusive in insects.

### Descending control of motor circuits

Explanations of how descending inputs from the brain control motor circuits have focused on two alternative modes of action: population coding versus command neurons^48,49^. The population model emphasizes the fact that a large number of descending neurons are often simultaneously active during a motor task^50^, and that each individual type of descending neuron may have a low-level, highly localized effect on a small number of muscles^51^. In this model, coordinated changes in motor output reflect the collective action of many different descending neurons, and only certain patterns of activity in these descending neurons - a population code - are capable of eliciting a behaviorally meaningful motor output. The command neuron model, in contrast, notes the profound and coordinated motor responses that can sometimes result from the activation of just a single class of descending neuron^52^. Because the motor responses elicited by a single command-type neuron can involve the coordinated activation of many different muscles, command-type descending neurons are generally thought to implement a relatively high-level control of motor circuits.

Computational models of backward walking in insects have posited the existence of bistable control switch for leg motor circuits, the setting of which determines whether motor rhythms are generated in a pattern that produces forward or backward walking^36,37^. Because this switch coordinately effects multiple motor outputs, it represents a high-level control embedded within the leg motor circuits. If such high-level directional switches exist in the *Drosophila* leg neuropils, then MDN, as a command-type descending neuron, could act by flipping these switches into the “backward” mode. Our data suggest however that this is not the case. We found that, of the two MDN-output neurons we examined in detail, each controls a discrete element of backward walking: hindleg tibia flexion during stance (LIN156) and leg lifting to initiate swing (LIN128). Many MDN-output neurons remain to be characterised, and it is possible that some of these may yet be found to be components of a high-level circuit switch coordinately effecting multiple motor features. Our data suggest however that at least some of MDN’s control is exerted at a lower level, close to motor output, and in a highly localized manner. The neuronal networks that control walking evidently operate in at least two stable states - one producing forward walking, the other backward walking. We propose that MDN may act through a distributed set of outputs, rather than a central control switch, to flip network dynamics into the backward state. In this model, MDN functions as a command-type neuron not because it acts at a higher level in motor control, but because the many discrete low-level changes it effects alter motor circuit dynamics so as to produce a clear and coordinated transition to backward walking.

Accordingly, we suggest that, in both the command-type and population coding scenarios, descending neurons can act in a highly localized, modulatory fashion, exerting their influence at the level of single muscles or muscle groups. In this view, it is not difficult to envision how evolution could shift the descending control of motor circuits up or down the continuum from a single-unit command-type control to a more combinatorial population-based control. Duplication of descending neurons and diversification of their outputs would result in a shift towards a more distributed control system, which might be favored under conditions in which flexibility or precision are advantageous. Conversely, a single descending neuron that initially has a highly specific and localized impact may become more potent over the course of evolution as it acquires new connections. An evolutionary shift in this direction might be favored if efficiency and reliability are more important than flexibility and precision.

In conclusion, we propose that there is no fundamental distinction between a command type of control and a population type of control. The former does not necessarily impinge upon motor circuits at a higher level than the latter. Rather, we suggest that these two opposing models are merely extremes on a continuum, and that both can be implemented through the distributed, low-level modulation of motor circuits. Where any single class of descending neuron falls on this spectrum depends, proximally, on the extent to which the many local modulations sum to produce a coordinated and behaviorally-relevant motor response, and ultimately, on the evolutionary trade-off between flexibility and efficiency.

## Acknowledgements

We thank the Janelia FlyLight, Fly Facility, Project Technical Resources teams, QBI infrastructure, microscopy, and IT teams and Samuel Kelly for technical assistance, Allan Wong for technical advice, and the Bloomington Drosophila Stock Center, Gerry Rubin, Vivek Jayaraman, David Anderson, Julie Simpson, Adam Claridge-Chang, Gilad Barnea, and Richard Mann for fly strains. We thank Josh Lillvis and other members of the Dickson group at Janelia and the van Swinderen group at QBI for comments on the manuscript, and Bruno van Swinderen at QBI for co-mentorship of K.F. This work was supported by funding from the Howard Hughes Medical Institute, the NHMRC, the ARC, and a University of Queensland Major Equipment and Infrastructure grant.

## Author Contributions

K.F. and B.J.D. conceived the study and wrote the manuscript, with contributions from all other authors. R.S. performed some of the fly-on-ball experiments, R. M. generated the split-GAL4 lines, M.D. and T.B. designed and manufactured core parts for the setup of high-speed video tracking of flies on ball under the supervision of A.B. K.F. performed all other experiments and analysed all the data.

## Extended Data Figures and Figure Legends

**Extended Data Fig. 1.**
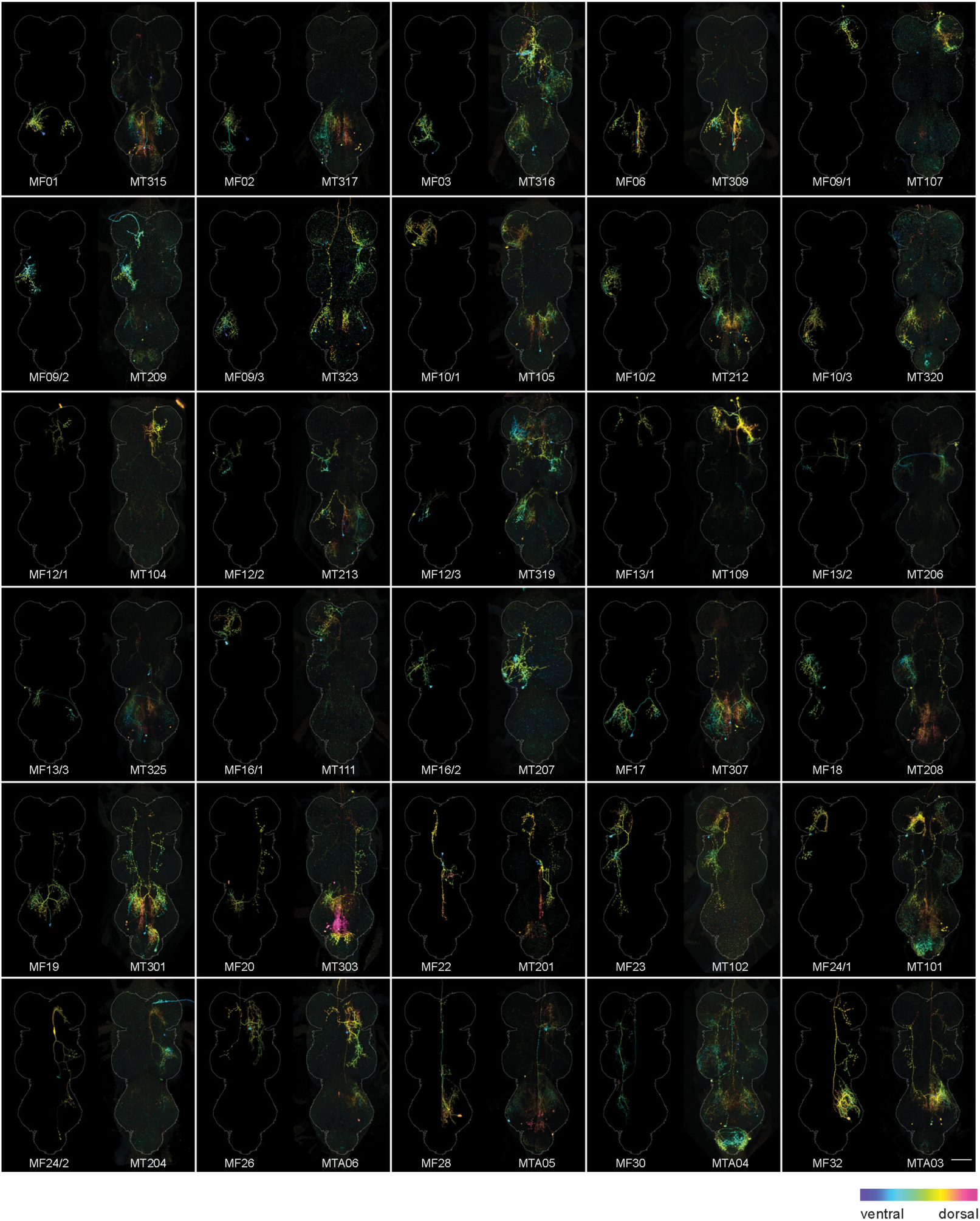
Comparison of cell types identified by trans-Tango and functional imaging. Cell types identified by both trans-Tango and functional imaging. All images are colorMIPs of registered confocal images. For each pair, the left image is an image of an MDN-responsive cell, segmented from an MCFO image obtained using the corresponding GAL4 driver. The right images is an unsegmented stochastic trans-Tango sample that appears to include the same cell type. Scale bar: 50 µm.

**Extended Data Fig. 2.**
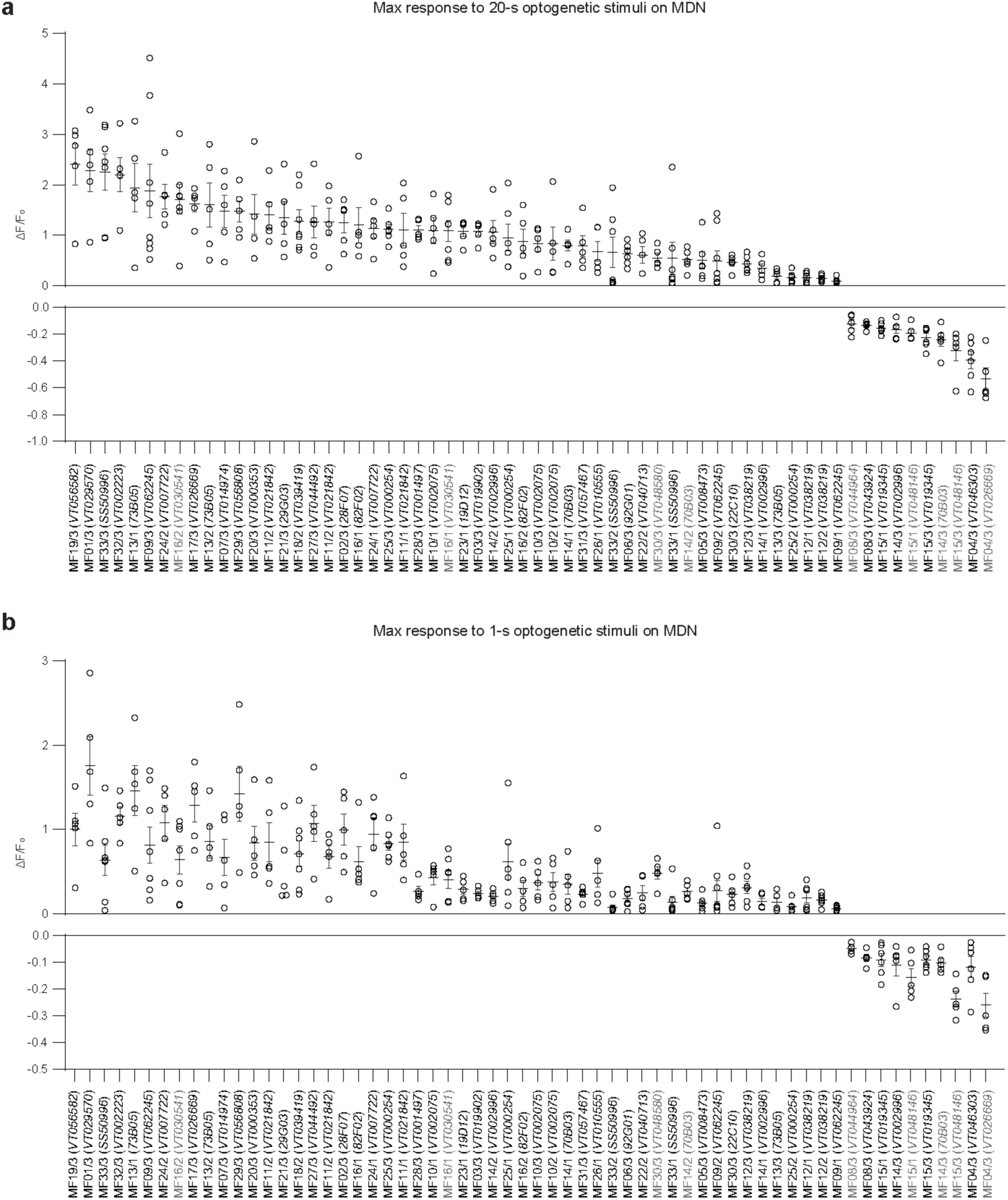
Calcium response profiles of MF cell types. Maximal ΔF/F responses of each cell upon (a) 20-s or (b) 1-s photoactivation of MDNs. Each data point (circle) represents a single fly, and bars show mean ±s.e.m. Samples are ordered in both plots by their response to the 20-s stimulus. “/1”, “/2”, and “/3” suffixes indicate T1, T2, and T3 cells, respectively. For some cell types, two independent driver lines were used. Black font indicates lines shown in Fig. 3.

**Extended Data Fig. 3.**
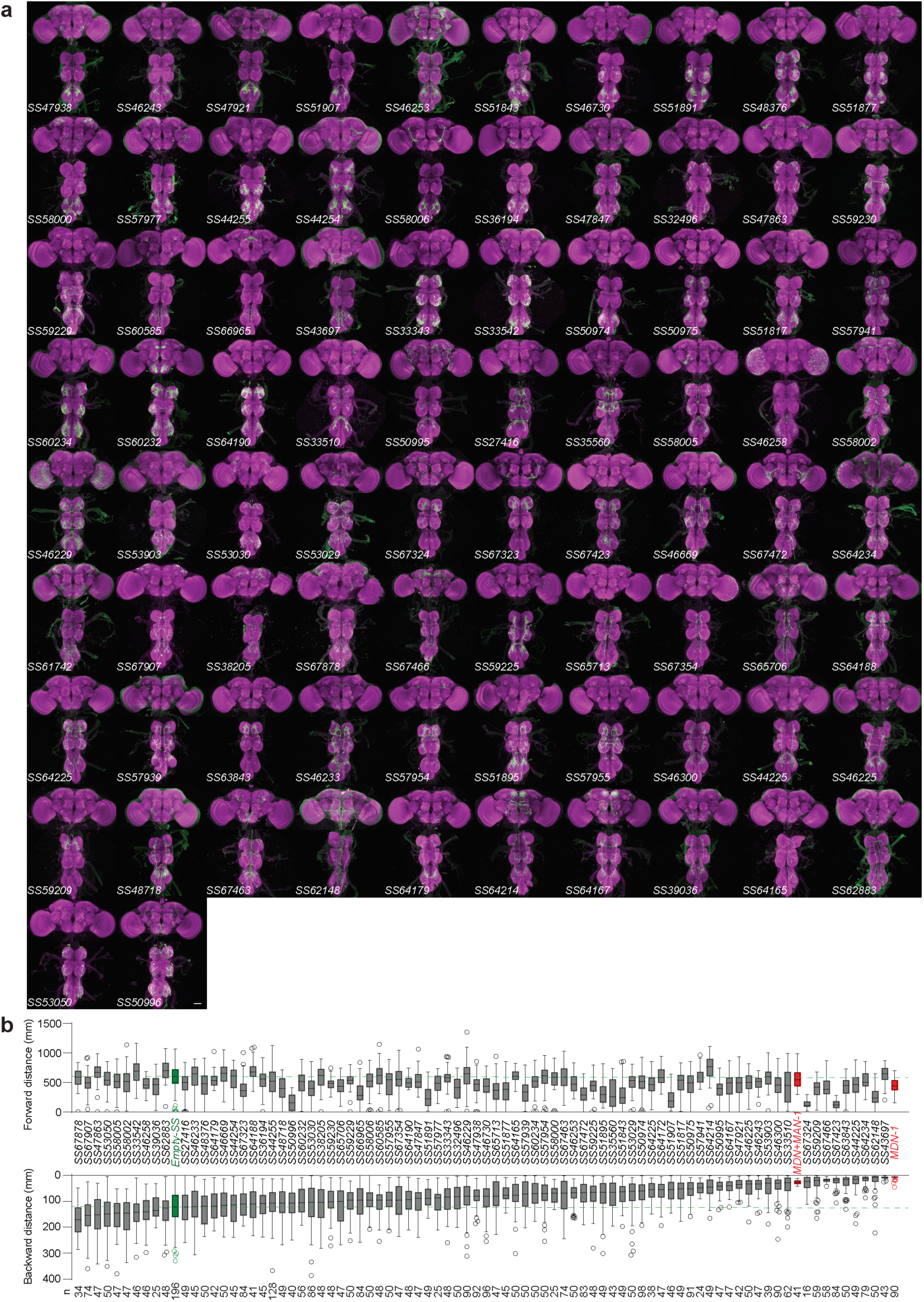
Anatomical and functional analysis of split-Gal4 lines labelling MDN-downstream neurons. a, Confocal images of central nervous systems for each split-GAL4 line shown in Fig. 4a, stained to reveal all synapses (nc82, magenta) and the targeted neurons (CsChrimson-mVenus, detected using anti-GFP, green). Scale bar: 50 µm. b, Box-and-whisker plots for the forward walking distance during 45 s without optogenetic stimulation (top), and total backward walking distance for the same flies during 9 successive 5-s red light pulses. Boxes show median and interquartile ranges. Bars extend down to the larger of the minimum value or the 25 percentile minus 1.5 times the interquartile range, and up to the smaller of the maximum value or the 75 percentile plus 1.5 times the interquartile range. Outliers are shown as circles. Positive and negative control lines are labelled in red and green, respectively. Green dashed lines show the median for the negative control, for comparison. Lines are ordered by the median backward walking distance.

**Extended Data Fig. 4.**
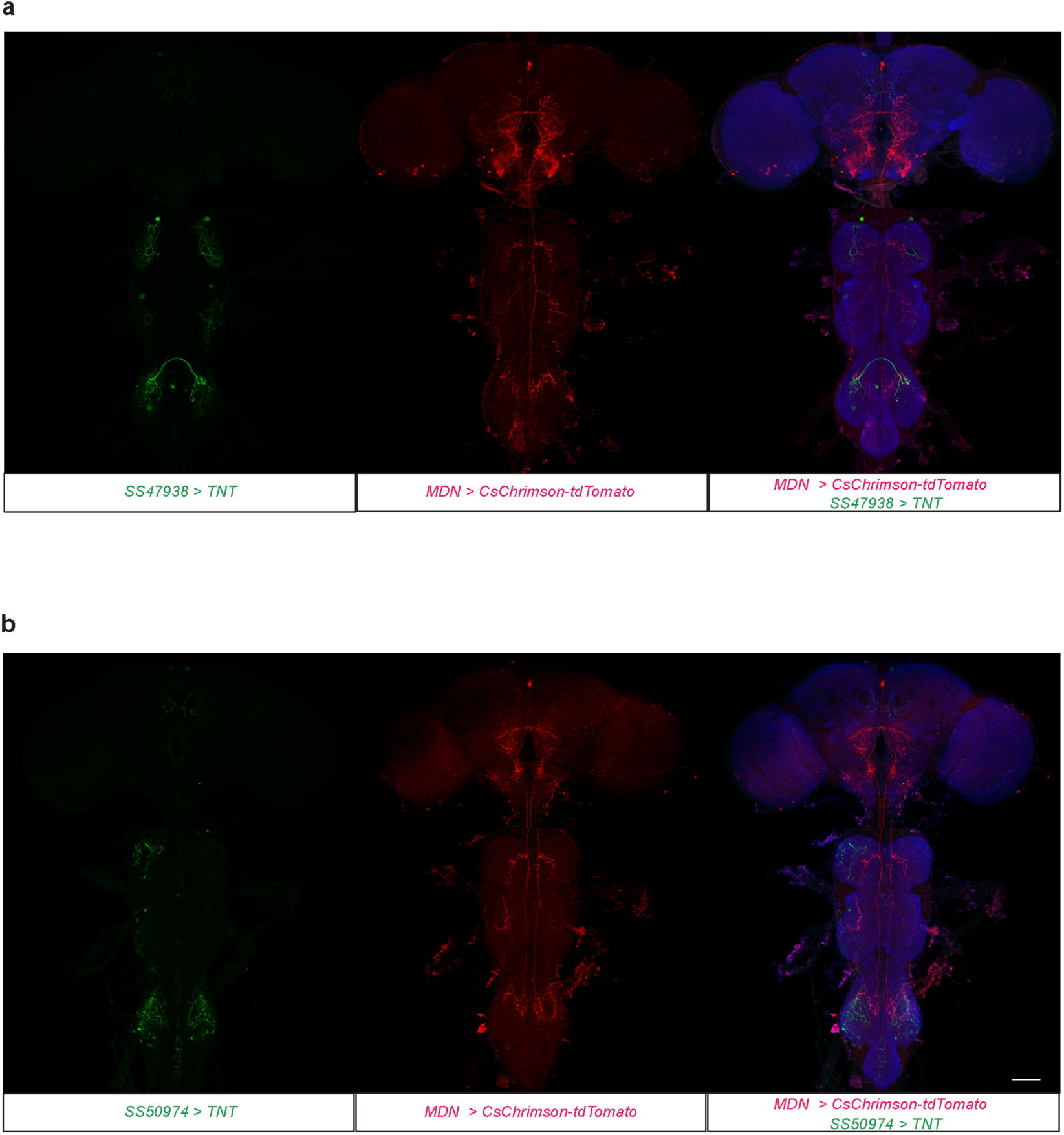
Expression of CsChrimson and TNT in flies with two split drivers. Confocal images of the central nervous systems of flies in which CsChrimson was expressed in MDN using a split-LexA driver and TNT was expressed in either LIN156 (a) or LIN128 (b) using a split-GAL4 driver. Samples were stained with anti-TNT (green), anti-RFP (to visualize CsChrimson-tdTomato, red) and nc82 (all synapses, blue). Scale bar: 50 µm.

**Extended Data Fig. 5.**
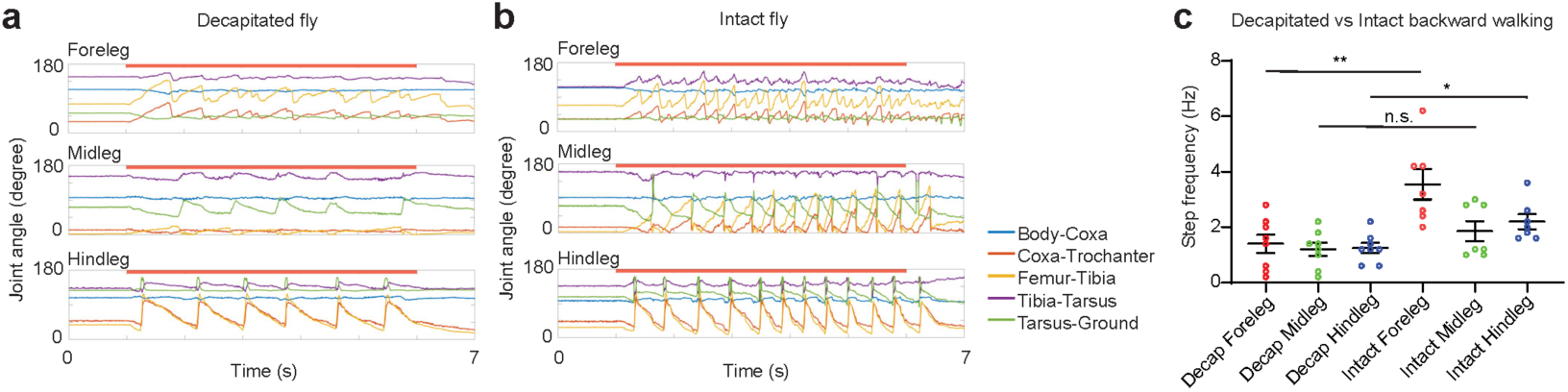
The backward walking program is largely intact in decapitated flies. a and b, Representative time series of joint angles in a decapitated (a) and intact (b) *MDN>CsChrimson* fly. Red bars indicate the 5-s red light stimulation. c, Backward step frequency for each leg in decapitated and intact flies. *N* = 7 flies for each group. Error bars show mean ± s.e.m.. **, *P*<0.01; *, *P*<0.05; n.s., *P*≥0.05; unpaired t-tests, two-tailed *P* values.

**Extended Data Fig. 6.**
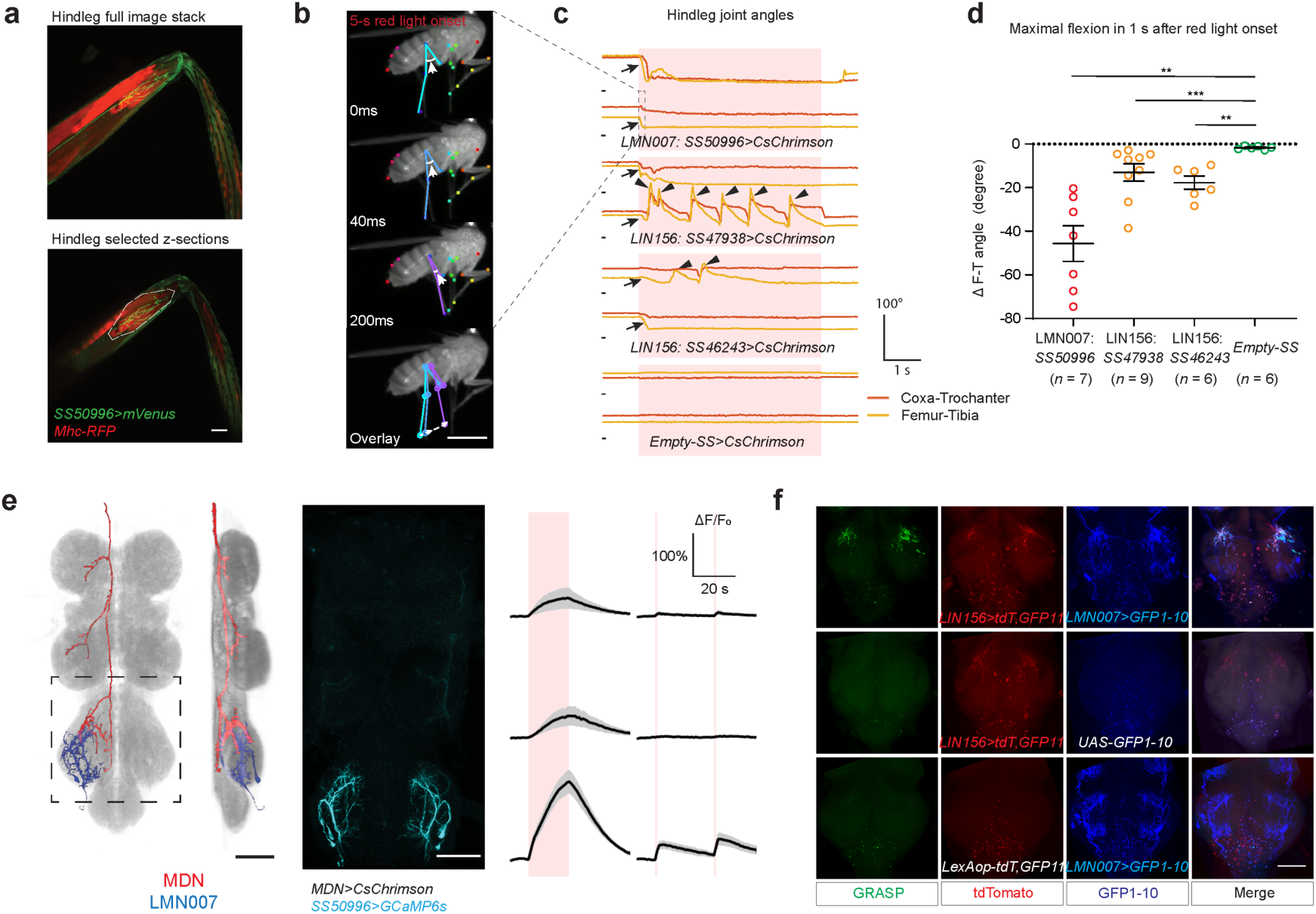
LIN156 triggers tibia flexion through LMN007. a, Top, maximum intensity projection of live fluorescence image of a hindleg, showing LMN007 innervation (mVenus, green) and all leg muscles (RFP, red). Bottom, selected z-slices highlighting innervation of the tibia reductor muscle (dashed line). b, Selected frames from a representative video of LMN007 activation in a suspended, decapitated fly, during a 5-s red light stimulus from t=0 ms. Arrows indicate the femur-tibia angles. Dashed line with arrows in the overlay image indicates sequence of movement. Flies were suspended rather than placed on a ball because simultaneous activation of foreleg and hindleg tibia flexor muscles in a fly on a ball would generate opposing pulling forces, potentially resulting in little or no movement of either joint. Scale bar: 1 mm. c, Hindleg joint angle time series, showing two representative traces for each genotype. Red shade indicates a 5-s pulse of red light. A short black line marks the origin for each trace (t=0 s, 0 degrees). Arrows indicate the initial femur-tibia flexion; arrowheads indicate extension events. The second set of traces is from the video shown in (b). d, Maximal femur-tibia joint angle change within 1 s after red light onset. ***, *P*<0.001; **, *P*<0.01, Mann-Whitney test, two-tailed *P* values. e, Left, registered segmented image of MDN (red), LMN007 (blue), and all synapses (gray, nc82). Scale bar: 50 μm. Middle, activated voxels (cyan) from calcium imaging of LMN007 upon MDN activation. Right, averaged responses for LMN007 to MDN activation (mean ± s.e.m., *N* = 7 flies), upon either a single 20-s stimulus or two 1-s light pulses (red shading). f, Live GFP fluorescence images showing reconstituted GFP (green) at likely synaptic connections between LIN156 (red, tdTomato live fluorescence, co-expressed with GFP11) and LMN007 (blue, anti-GFP staining against GFP1-10). The imaged areas correspond to the dashed box in e. Scale bar: 50 μm.

**Extended Data Fig. 7.**
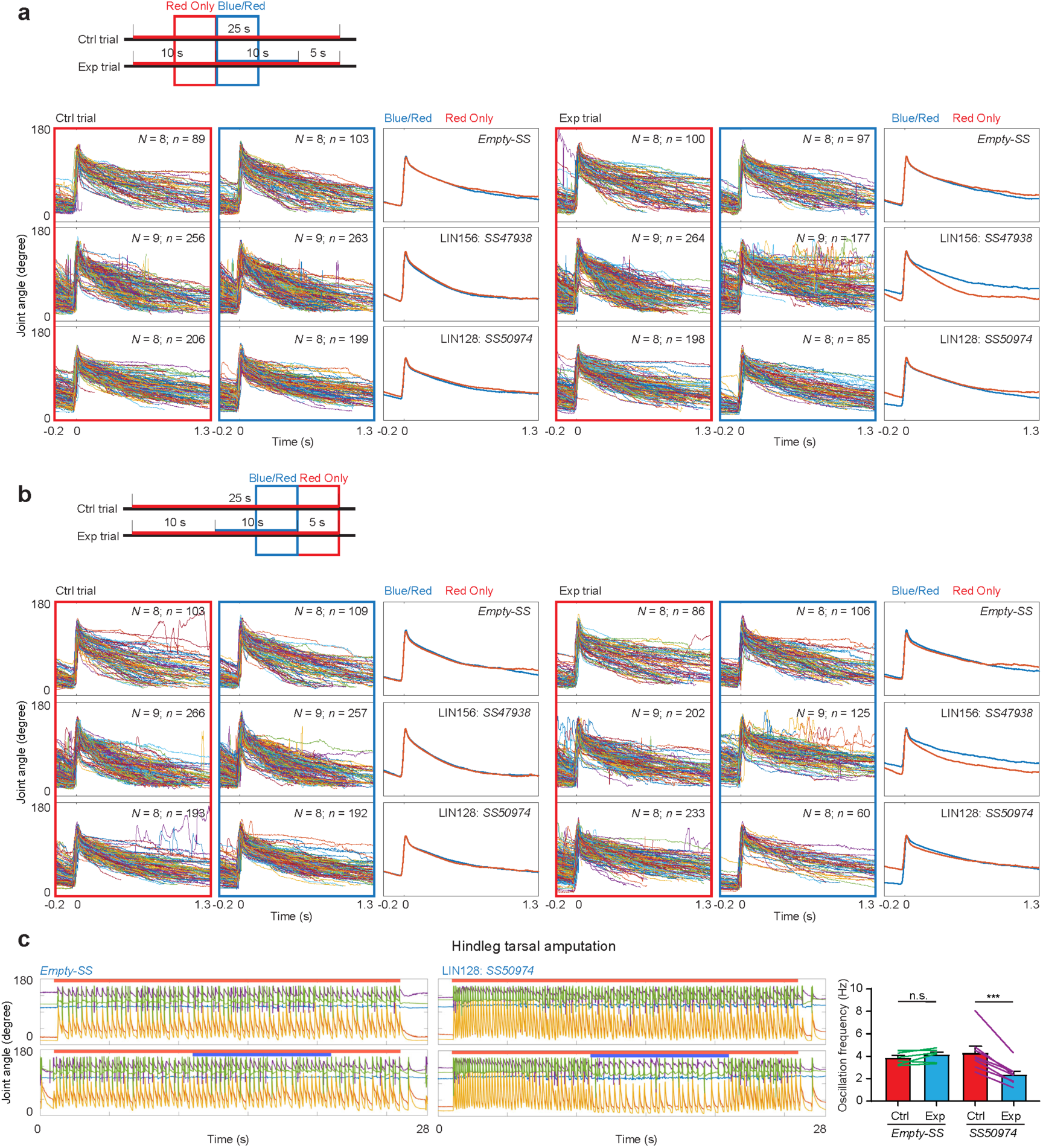
Acute silencing of LIN156 or LIN128 leads to distinct defects in backward walking. a and b, Overlaid and average time series of femur-tibia joint angle for all the steps extracted from the indicated time windows. The same dataset as shown in Fig. 7 were analysed using within-trial comparisons. c, Left, representative time series of joint angles in hindlegs amputated at tarsus. Red bar indicates red-light stimulus; blue bar indicates blue-light stimulus. Right, quantification of oscillation frequency of amputated hindlegs. *N* = *7* flies for *Empty-SS* and *N* = 8 for *SS50974*. Error bars show mean ± s.e.m. ***, *P*<0.001; n.s., *P*≥0.05; paired t-tests, two-tailed *P* values.

## Supplementary Information

### Supplementary Videos

**Supplementary Video 1 | Joint kinematics during forward and backward walking.**

A tethered *MDN-1-GAL4>CsChrimson* fly spontaneously walking forward on a ball, then walking backwards upon presentation of a 5-s red-light stimulus (red square). Leg segments (blue) and joints and tips (red) on one side of the body were labelled using DeepLabCut. The trajectory of tarsal tips is plotted in white for forward walking and in red for backward walking. Recorded at 200 fps and rendered at 40 fps.

**Supplementary Video 2 | Amputation experiments reveal a dominant role for the hindlegs in backward walking.**

*MDN-1-GAL4>CsChrimson* flies walking on a ball, before, during, and after a 5-s red-light stimulus (red squares). Flies shown from left to right have the fore-, mid- or hindlegs bilaterally amputated at tarsus, respectively. Recorded at 200 fps and rendered at 40 fps.

**Supplementary Video 3 | LIN156 activation triggers tibia flexion.**

A decapitated *SS47938>CsChrimson* fly presented with five 50-ms red light pulses (red squares) at 1-s intervals. All presentations elicit tibia flexion; for the third and the fifth stimuli, flexion is followed by re-extension. Recorded at 200 fps and rendered at 40 fps.

**Supplementary Video 4 | LIN128 activation triggers leg lifting.**

A decapitated *SS50974>CsChrimson* fly presented with five 5-ms red light pulses (red squares) at 1-s intervals. All presentations elicit hindleg lifting. Recorded at 200 fps and rendered at 40 fps.

**Supplementary Video 5 | Coordinated stepping induced by LIN128 activation.**

A decapitated *SS50975>CsChrimson* fly presented with a 5-s red-light pulse (red square), resulting in coordinated stepping across all 6 legs. Recorded at 200 fps and rendered at 40 fps.

**Supplementary Video 6 | Silencing LIN156 during backward walking results in a slower stroke.**

A decapitated *MDN >CsChrimson, SS40738>GtACR2* fly presented with a 25-s red-light pulse (red square) to activate MDN and trigger backward walking, during which a 10-s blue light pulse (blue square) was presented to silence LIN156 neurons. Flexion of the hindleg femur-tibia joint is noticeably slower during the blue light stimulus. Recorded and rendered at 200 fps.

**Supplementary Video 7 | Silencing LIN128 during backward walking delays swing phase.**

A decapitated *MDN >CsChrimson, SS50974>GtACR2* fly presented with a 25-s red-light pulse (red square) to activate MDN and trigger backward walking, during which a 10-s blue light pulse (blue square) was presented to silence LIN128 neurons. The hindleg femur-tibia joint reaches an abnormally acute angle and swing phase is noticeably delayed during the blue light stimulus. Recorded and rendered at 200 fps.

### Supplementary Table

**Supplementary Table 1.**
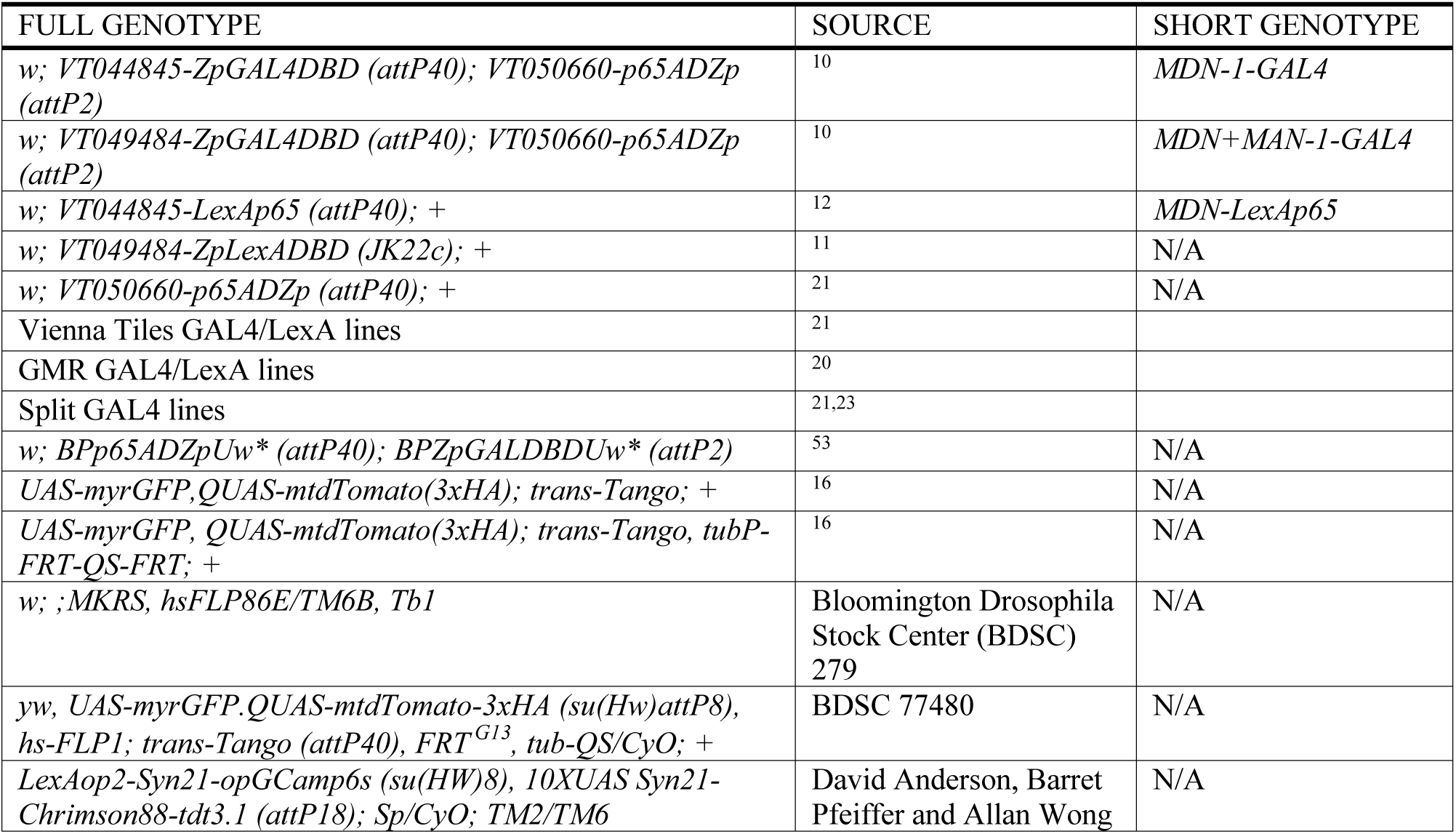
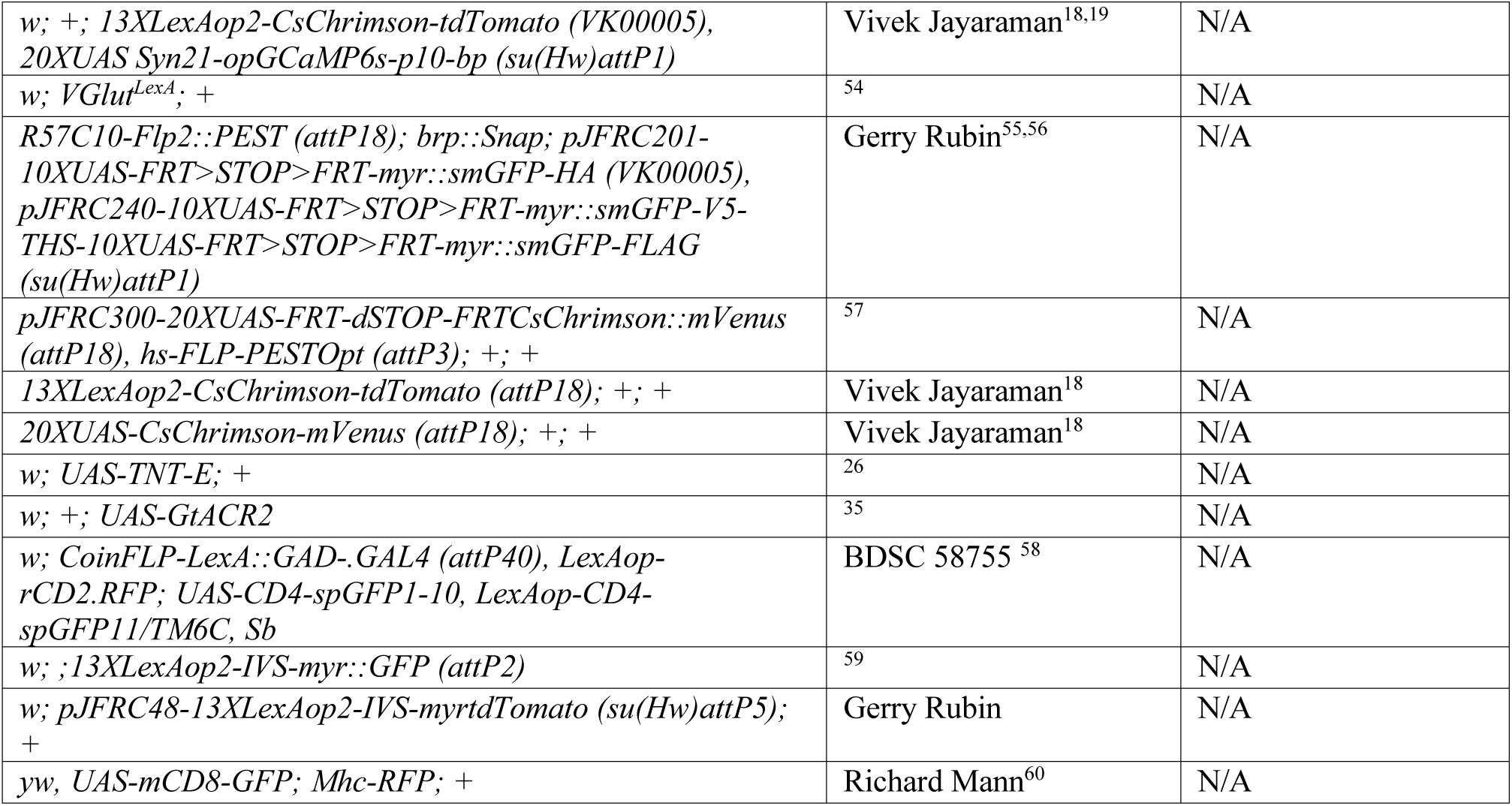
Fly Stocks.

## Online Methods

### Fly Stocks

See Supplementary Table 1 for genotypes of *Drosophila* used in this study. All flies were raised on standard semi-defined media at 25°C, 50% relative humidity and 12hr-12hr light cycle except for flies used in the trans-Tango dataset, which were raised at 18 °C. All flies for behavioral experiments or functional imaging were sexed within 2 days after eclosion and aged on fresh food supplemented with all-trans retinal (0.2 mM) and kept in the dark (food tube wrapped with foil). All flies for immunohistochemistry were sexed within 2 days after eclosion and aged on fresh food. Both males and females were used in Fly Bowl experiments, CsChrimson activation experiments using tethered flies, and immunohistochemistry. For calcium imaging, MDN amputation and GtACR2 neuronal epistasis experiments, only females were used. For all behavioral experiments and immunostainings except for trans-Tango, 3-8 day old adult flies were used. For calcium imaging, 3-20 day old adult flies were used. For trans-Tango experiments, 15-40 day old flies (raised at 18°C) were used.

### Behavioral Assays

#### Fly Bowl

Neuronal epistasis experiments in Fig. 4b and Extended Data Fig. 3b were conducted as in ref. 12 using the Fly Bowl setup^25^. Briefly, video recordings were performed under backlit infrared LEDs at 30 fps with a resolution of 1024 × 1024 pixels. A group of ∼25 flies were loaded into the bowl-shaped arena and allowed to walk for 60 s before the first optogenetic simulation. The first 45 s of this period were used to compute the spontaneous forward walking speed in Extended Data Fig. 3b. A red light source at 627 nm wavelength illuminated the entire arena uniformly to activate CsChrimson. Each red light stimulus was 5 s in duration and in 60 s intervals (onset to onset). In total 9 episodes of red light stimuli were applied in one experiment with 3 consecutive episodes for each of 3 intensity levels from low to high: 0.158 mW/mm^2^, 0.288 mW/mm^2^ and 0.456 mW/mm^2^.

#### High-speed videos of tethered flies

Flies were anaesthetized using a cold block set to 0-1 °C and placed in a fly-shaped groove. A drop of adhesive “Not-A-Glue” (Bondic, Canada) was applied to the notum of the fly using a thin metal wire. A bent metal pin fixed on one end of a rod was lowered by a manual manipulator to gently touch the adhesive. Then the adhesive was cured by LED light (provided with the adhesive). When decapitated flies were used, we cut off fly’s head with fine scissors (Fine Science Tools, CA) before gluing. The tethering process normally takes less than 3 minutes. The tethered flies were then given a styroform ball (∼5 mm diameter) to hold before experiments to keep them from struggling. The rod with fly tethered is then attached on a manual manipulator in the ball setup. The compressed air supporting the ball was carefully adjusted to the minimal level that can spin the polypropylene ball (PP ball, 6 mm in diameter, Spherotech GmbH, Germany) without the fly stepping on it. The fly was then moved carefully to the top of the PP ball supported by a custom-built ball holder (Electronics workshop, Zoological Institute, University of Cologne). A top camera was used to monitor the fly’s position on ball so that the fly can be properly centred. From the side view, we used the minimal projected area of the wings as a sign to adjust the roll of the fly. A custom-built infrared-LED ring (wavelength: 880 nm, Electronics workshop, Zoological Institute, University of Cologne) was positioned around the camera lens for illumination. A few initial videos were acquired at 200 fps with a resolution of 464 × 464 pixels from a CameraLink camera VC-2MC-M340 (Vieworks, Korea) equipped with a telecentric lens TEC-M55 (Computar, USA). For majority of the experiments, a USB3 camera GS3-U3-23S6M-C (FLIR, Canada) equipped with a 75 mm zoom lens with extension tube (to further increase the zoom so that the field of view on the 1/1.2 inch CMOS chip is ∼12 mm in width) and an infrared filter was controlled by Spinnaker software (FLIR, Canada) to acquire videos from the side view of fly. For the FLIR camera, videos were initially acquired at 200 fps at a resolution of 960 × 800 and then cropped and downsized to 464 × 464 to match the videos taken from the Vieworks camera. Data from both Vieworks and FLIR cameras were analysed by the same DeepLabCut model. Only for *SS50975* activation experiments, two Basler acA1920-155um cameras (Basler AG, Germany) equipped with Infinistix 1X 94 mm lens (with a 5.8 mm aperture retainer ring and an infrared filter; Infinity Photo-optical Company, CO) were used under control of Pylon software (Basler AG, Germany) to take videos simultaneously from both sides of the body. In those experiments, frame rate was 200 fps and frame size was 960 × 480. Videos from Basler cameras were manually scored. An Arduino UNO (Arduino.cc) was used to synchronize cameras and optogenetic stimuli. A custom built 660 nm diode laser with fibre coupling (Electronics workshop, Zoological Institute, University of Cologne) was used to deliver the red light stimulus for CsChrimson. A fiber coupled 470 nm LED (M470F1, Thorlabs, NJ) controlled by a custom LED driver (QBI Workshop, Australia) was used to deliver the blue light stimulus for GtACR2. Multimode optical fibers (Ø200 µm, 0.39 NA for red light; Ø200 µm, 0.50 NA for blue light; Thorlabs, NJ) were used to deliver light to the flies (both with SMA coupling to source and clean-cut bare end pointing to sample). The red light intensity used for 50-ms (Fig. 5f,g and Supplementary Video 3), 5-s (Fig. 1, Fig. 5c-e, Fig. 6d,e,g,h, Extended Data Fig. 5, Extended Data Fig. 6c,d, and Supplementary Videos 1,2,5) and 25-s pulses (Fig. 7, Extended Data Fig. 7, and Supplementary Videos 6,7) was approximately 0.73 mW/mm^2^ and the intensity for 5-ms pulses (Fig. 6c, Fig. 6f and Supplementary Video 4) was approximately 3.8 mW/mm^2^. The blue light intensity (Fig. 7, Extended Data Fig. 7, and Supplementary Videos 6,7) was approximately 0.11 mW/mm^2^. For *MDN>CsChrimson* activation experiments, we first allowed the flies to walk freely in order to capture spontaneous forward walking episodes and then triggered backward walking using a 5-s constant red light stimulus. For tarsal amputation experiments, we first recorded from intact flies and then cut off the tarsal tip at the middle of the tarsus on cold block with a pair of fine scissors before recording again from the same flies with identical protocol. For *SS>GtACR2* neuronal epistasis experiments, we first tested if a decapitated fly could walk backwards steadily during the 25-s red light stimulus. Only flies that showed stable baseline backward walking throughout the 25-s window were further tested. We performed exactly the same number of control trials and experimental trials in an alternating sequence on each fly.

### Immunohistochemistry

Immunohistochemistry was performed on the central nervous system (CNS) dissected from adult flies, following protocols in https://www.janelia.org/project-team/flylight/protocols. Samples for MCFO, mVenus and GFP staining and tdTomato/TNT double staining were DPX mounted. Samples for trans-Tango and GRASP experiments were mounted in Vectashield (Vector Laboratories, CA). Confocal stacks were obtained at 1 µm intervals using an LSM 710 microscope (Carl Zeiss AG, Germany) with a 20X objective, or a 40X objective for MCFOs.

For trans-Tango experiments, the flies were aged at 18 °C for 15-40 days before dissection. We found in pilot experiments that *MDN-1>trans-Tango* flies showed only sparse or no labelling of post-synaptic cells, but a *UAS-CsChrimson-mVenus* transgene enhanced the trans-Tango signal dramatically even in the absence of a red light stimulus or retinal food. For stochastic trans-Tango, two genotypes with different *hs-FLP* transgenes were used and the data pooled. Flies were heat-shocked at 37 °C for 1-3 sessions each lasting 2 hours at various stages from first-instar larvae to mid-pupae. We diversified the heat-shock protocol in order to get sparse labelling as well as high coverage of post-synaptic cells. When multiple heat shocks were performed, they were spaced by at least 1 day. More than a thousand samples were dissected and stained, of which 541 samples with sparse to medium expression were registered against a VNC template (JRC2017,ref. 17) and further analysed. The antibodies and concentrations used in those experiments were as follows: primary rabbit anti-GFP (Merck/Millipore AB-3080P, 1:1000), primary rat anti-HA (Roche 11867423001, 1:100), primary mouse anti-Bruchpilot (Developmental Studies Hybridoma Bank nc82, 1:20), secondary Alexa Fluor 488 goat anti-rabbit (Thermo Fisher Scientific A-11034, 1:500), secondary Alexa Fluor 555 goat anti-rat (Thermo Fisher Scientific A-21434, 1:800) and secondary Alexa Fluor 647 goat anti-mouse (Thermo Fisher Scientific A-21236, 1:500).

For mVenus and GFP staining, the antibodies used were as follows: primary rabbit anti-GFP (Thermo Fisher Scientific A-11122, 1:1000), primary mouse anti-Bruchpilot (1:30), secondary Alexa Fluor 488 goat anti-rabbit (1:800) and secondary Alexa Fluor 568 goat anti-mouse (Thermo Fisher Scientific A-11031, 1:400). For MCFO, the majority of experiments were hybrid immunohistochemistry/chemical tag stainings, following the protocol listed in the abovementioned website (see also ref. 61). The antibodies and dyes used were Cy2 SNAP-tag ligand (Luke Lavis, JRC, 10 μL/mL for final concentration of 2 μM), primary rat anti-FLAG Tag (Novus Biologicals NBP1-06712, 1:200), primary rabbit anti-HA Tag (Cell Signal Technologies 3724S, 1:300), secondary ATTO647N goat anti-rat (Rockland 612-156-120, 1:300), secondary AF594 donkey anti-rabbit (Jackson ImmunoResearch 711-585-152, 1:500) and DL550 mouse anti-V5 (AbD Serotec MCA1360D550GA, 1:500). A few experiments were immunohistochemistry only and only differed from the hybrid protocol in that mouse anti-Bruchpilot (1:30) and Cy2 Goat anti-mouse (Jackson ImmunoResearch 711-585-152, 1:600) were used instead of the Cy2 SNAP-tag ligand. For tdTomato and TNT double labelling, the antibodies used were as follows: primary rabbit anti-TNT (Statens Serum Institute 65873, 1:3000), primary rat anti-RFP (Chromotek 5f8-20, 1:500), primary mouse anti-Bruchpilot (1:30), secondary Alexa Fluor 488 goat anti-rabbit (Thermo Fisher Scientific A-11034, 1:800), secondary Cy3 goat anti-rat (Jackson ImmunoResearch 112-165-167, 1:1000) and secondary Cy5 goat anti-mouse (Jackson ImmunoResearch 115-175-166, 1:1000). For GRASP experiments, we used polyclonal antibody rabbit anti-GFP (AB-3080P, 1:1000) followed by secondary Alexa Fluor 647 goat anti-rabbit (Thermo Fisher Scientific A-21245, 1:500) to label GFP1-10 and imaged the live fluorescence for reconstituted GFP and tdTomato, which was co-expressed with GFP11. For leg muscle innervation in Extended Data Fig. 6a, the flies were fixed in 4% paraformaldehyde overnight at 4°C. After 5 times of 15-minute washing in 0.3% PBST, we removed the legs with forceps and mounted them in Vectashield. We then imaged live fluorescence of mVenus and RFP for nerves and muscles respectively.

### Calcium imaging

All calcium imaging experiments were performed on *ex vivo* preparations of the fly CNS. The entire CNS of female flies were dissected in extracellular solution comprising (in millimoles): 103 NaCl, 3 KCl, 5 N-Tris(hydroxymethyl)methyl-2-aminoethanesulfonic acid (TES), 8 trehalose, 10 glucose, 26 NaHCO3, 1 NaH2PO4, 2 CaCl2, and 4 MgCl2 (pH near 7.3 when bubbled with 95% (vol/vol) O_2_ and 5% (vol/vol) CO_2_ (carbogen)). The extracellular solution were bubbled with carbogen and pre-chilled with ice before dissection. After dissection, the tissues were quickly transferred to a custom chamber filled with fresh extracellular solution and stuck to a fresh glass coverslip placed at the bottom of the chamber. The chamber was then placed on the stage under microscope and constantly perfused with fresh extracellular solution bubbled with carbogen throughout the experiments. The room temperature was controlled at 23°C.

Two-photon imaging was performed on a Thorlabs Bergamo II microscope equipped with a Galvo-Resonant scanner (Thorlabs, NJ). A piezo objective focus module with 400 µm travel distance (Physik Instrumente GmbH & Co. KG, Germany) was used to control an Olympus XLUMPLFLN 20X water immersion objective (Olympus, Japan). A Mai Tai DeepSee Ti:Sapphire laser (Spectra-Physics, CA) was tuned to 920 nm for two-photon stimulation. A Pockels Cell (Conoptics Inc., CT) was used to control the laser intensity and ramp it while scanning deeper into the tissue. A 617 nm LED source (M617F1, Thorlabs, NJ) controlled by a LED driver (DC4104, Thorlabs, NJ) was coupled to the two-photon light path to deliver the optogenetic stimuli through the objective. A 594 nm long pass dichroic was used pass the 920 nm laser and 617 nm red light and reflect fluorescence from the sample to a pair of GaAsP PMTs (Hamamatsu, Japan). A 562 nm long pass dichroic was placed between the two PMTs and a 525/50 nm band pass filter was placed in front of the PMT that was used to detect light from GCaMP. During an imaging session, an Arduino UNO (Arduino.cc) was used to trigger and synchronize two-photon imaging and optogenetic stimulus.

All imaging experiments were performed under the streaming mode in Thorimage software (Thorlabs, NJ). We scanned 27 effective slices spaced by 6 µm plus 3 flyback slices (discarded during analysis) to cover a whole VNC volume. The imaging frame was set to 512 × 1024 pixels and zoomed to just cover the full VNC. Under such a condition, the imaging speed was approximately 30 frames or 1 VNC volume per second. The laser power on sample was ranging from 1.5 mW to 20 mW dependent on the driver line’s strength and depth of scanning plane. We performed recording sessions containing 60 volumes (∼1 minute) and gave the sample either 1 red light stimulus lasting 20 s (onset from imaging start: 10 s; see Fig. 3d-f, Fig. 5a, Fig. 6a, Extended Data Fig. 2a, and Extended Data Fig. 6a,e) or 2 stimuli each lasting 1 s (onset from imaging start: 10 s and 40 sec; see Fig. 5i, Extended Data Fig. 2b, and Extended Data Fig. 6e). Each red light stimulus contains a train of 5 ms pulses at 50 Hz. The light power (not peak power; 25% duty cycle as a result of 5 ms pulses at 50 Hz) measured at the sample was 0.81 mW. Given the field of view under the 20X objective was approximately 0.9 mm in diameter, we calculated the light intensity was approximately 1.27 mW/mm2. In a typical experiment, we repeat 10 such 60-s sessions on each sample, with a 15-s interval between sessions.

### Functional imaging screen

Starting from relatively broad lines and patterns we saw from trans-Tango experiments, we selected ∼300 GAL4 or LexA lines from the GMR^20^ and VT collections^21^ with progressively sparser expression patterns until we can pinpoint any single cell types from the calcium imaging patterns responding to MDN activation. We used *MDN-1-GAL4* (*VT044845*-*ZpGAL4DBD in attP40, VT050660-p65ADZp in attP2*, ref. 10) to drive expression of Chrimson88 specifically in MDNs when we imaged LexA lines. We used a split MDN-LexA (*VT049484-ZpLexADBD in JK22c, VT050660-p65ADZp in attP2*, ref. 11) to drive expression of CsChrimson in MDNs when we imaged GAL4 lines. The latter used the same enhancer combinations as the *MDN+MAN-1-GAL4*^10^ and expressed in both MDNs and MANs but no other central neurons. To exclude the possibility that MANs were responsible for activating a given cell type, we also used a MDN-LexA (*VT044845-LexA*, ref. 12), which is broader but does not label MAN, to confirm the response we saw from a positive GAL4 line is attributable to MDN activation. For the screen, we typically imaged 2 samples per line with the 20-s photo-stimulation protocol and rescreened the few lines that showed inconsistent patterns. Combining the results from both GAL4 and LexAs, we identified 33 cell types of which the calcium imaging patterns can be clearly matched to a neuron from MCFO patterns driven by lines containing the same enhancer as those we used for calcium imaging. Many cell types were hit by multiple GAL4 or LexA lines. We selected at least one line for each cell type and expanded the sample numbers to at least five, with which we applied both 20-s and 1-s stimulation protocols, as shown in Fig. 3 and Extended Data Fig. 2.

### Neuron segmentation

To segment an MF neuron type, we obtained stochastic labelling images using GAL4 lines containing the same enhancers that were used for calcium imaging to drive expression of a MCFO reporter ^22^. To segment an MT neuron type, we used images from the stochastic trans-Tango dataset in Fig. 1b. In both cases, we selected images that showed relatively sparse patterns surrounding the neuron of interest. For MDN (Fig. 5a, Fig. 6a and Extended Data Fig. 5e), we used an image from the MDN stochastic labelling dataset reported in ref. 12. All VNC images were non-rigidly registered^62^ to JRC2017 template^17^ and brains to JFRC2013 template^63^. Using the software VVD Viewer ^17,64,65^, we rendered the registered image stacks in 3D and manually masked other neurons co-labelled in the image and segmented out the neuron of interest.

### Generation of split-GAL4 lines targeting MF cell types

As part of a large effort to systematically targeting cell types in leg neuropils (R. M., K. F., and B. D, *in preparation*), we used images of targeted cell types to generate Color-MIP masks and search through the images of GMR and VT lines^17^ to obtain enhancers that may contain the cell type of interest. We made combinations of these enhancers pairwise in split-GAL4 halves^21,23^ and tested their expression patterns by immunostaining. For those pairs that sparsely labelled the neuron types of interest, we stabilized them by double balancing the AD and DBD halves to obtain the *SS* lines.

### Quantification And Statistical Analysis

#### Quantification of calcium imaging results

From the 4-D raw data of imaging, we first concatenated 10 of the 60-s recording sessions on the same sample to create a 600-frame hyperstack containing 1024 × 512 × 27 voxels. We then performed motion-correction on each of the 27 slices using the NoRMCorre algorithm^66^. We also generated two GCaMP6s kernels for both the 20-s and 1-s stimuli. To do this, we manually defined ROIs on neurons that showed strong activation with short delay and calculated the ΔF/F_0_. By averaging multiple such ROIs, we created smooth response curves that capture typical response dynamics to our MDN activation protocols. We then calculated the cross-covariance between the time series of each voxel and the corresponding kernel using the Matlab function “xcov” with the “maxlag” parameter set to 5. This parameter allowed us to detect time-shifted responses that could come from polysynaptic connections to MDN. The cross-covariance analysis resulted in 11 values for each voxel. We used the maximal positive values or 0 if all 11 were negative to reconstruct an image stack that represents the “activated” voxels (cyan in Fig. 3b,c, Fig. 5a,i, Fig. 6a and Extended Data Fig. 6e). Likewise we used the minimal negative values or 0 if all 11 were positive to reconstruct an image stack that represents the “inhibited” voxels (red in Fig. 3b). We compared these images with immunostainings of the same or other driver lines to identify neurons that respond to MDN activation or select sparser lines to narrow down the pattern. We also used these images as a guide to define ROIs on neurons of interest to calculate ΔF/F0. We manually defined ROIs using the freehand or polygon tools in Fiji^67^. For each sample, the ROI was defined on a single imaging plane that showed strong responses and large areas for the given neuron type. For neuron types containing multiple cells with overlapping neurites, the ROIs may cover multiple neurons of the same type. For neuron types containing a single neuron and the counterparts in each hemisphere do not overlap, we defined the ROIs on the neuron that showed stronger response in each sample. To calculate ΔF/F0, we took the averaged intensity in each ROI from frame 2-9 for each of the 60-s imaging sessions as the baseline F0. For each sample, we then calculated averaged ΔF/F0 of the 10 imaging sessions. The averaged ΔF/F0 from multiple samples were used to compute the response curves (see Fig. 3d-f, Fig. 5a,i, Fig. 6a, and Extended Data Fig. 6e) and quantify the maximal responses (see Extended Data Fig. 2) for each cell type.

#### Quantification of Fly Bowl assays

Videos recorded from the Fly Bowl were tracked and analysed using a pipeline based on Ctrax^68^. Briefly, all flies were first tracked in each frame throughout the video. Forward/backward speed (in Ctrax output reported as “du_ctr”) were further analysed to generate results in Fig. 4b and Extended Data Fig. 3b. This excluded the speed component of sideways walking (relative to the fly’s heading). Spontaneous forward walking and triggered backward walking distance for each fly was computed individually and results for all flies from multiple assays of each genotype were pooled. For forward walking distance, the total forward/backward distance for flies walked in the initial 45 s (15 s prior to the first episode of red light stimulus) of the assay was computed. For backward walking distance, the accumulative backward distance (a backward speed threshold >=1.5 mm/s was applied as in ref. 12) during the 9 episodes of red light stimuli (3 red light levels, 3 episodes each level, 5 s each episode, in total 45 s) was computed.

#### Joint kinematics

An artificial neural network was trained using DeepLabCut software^15,69^ to mark four joints and the tarsal tip of each leg, two antenna and three points on the abdomen (as illustrated in Fig. 1a) of a fly either walking on ball or being suspended. In total 780 frames out of 39 videos sized 464 by 464 were manually marked as the training set. The frames were carefully chosen to represent flies under different conditions and performing different motor tasks and a few pilot networks were trained and evaluated to extract some of the frames included in the final training set. The final network was trained by this training dataset from the default resnet101 weights for 1030000 iterations. All the videos analysed by the network were overlaid with the markings (see Fig. 1a) for manual quality control. Tracking errors were usually limited to very few frames. Time series of joint angles (as illustrated in Fig. 1b) and tarsal tip y positions were batch computed and plotted to facilitate human inspection. For the dataset in Fig. 1b-h, only videos that did not contain obvious tracking errors for any joints were used. For all cases in which femur-tibia and/or coxa-trochanter angles were plotted and quantified, manual quality control was applied on the relevant joints to exclude videos that had obvious mistracking. Z-scored tarsal tip y positions were then used to detect the peak of each swing phase (referred to as “swing peak” hereafter) by Matlab function ‘findpeaks’. For all cases where individual steps were extracted or stepping frequency were quantified, manual quality control was applied to correct the errors in swing peak detection.

For Fig. 1b-h, we used a dataset containing one episode of spontaneous forward walking and one episode of optogenetically triggered backward walking for each of the 7 flies. Manual proofreading ensured errors in joint position tracking and swing peak detection (see above) were minimal for this dataset. Using the swing peaks, we aligned all the steps across flies and averaged the tarsal positions and joint angles to plot the data in Fig. 1c,d. For Fig. 1d, the 0.5 s time window contained the swing peak at frame 50, 49 preceding frames, and 50 subsequent frames after it. If a neighbouring swing peak falls into this time window, we truncated the step. So for each step, the boundary was either the abovementioned time window or shortly after the previous swing peak or before the next swing peak. The grey areas indicating swing phase were determined by the inflection point of the positions of tarsal tips of the averaged steps. The step size was defined as the distance between Anterior Extreme Position (AEP) and Posterior Extreme Position (PEP). The swing stroke amplitude was defined as the distance between the Dorsal Extreme Position (DEP) and the line segment connecting AEP and PEP. The joint angle range was defined as the maximum minus the minimum of each joint angle during a step. To plot Fig. 1e-h, we calculated the mean of relevant parameters from all steps for each fly, which represented a single data point in these plots.

For Fig. 1i,j we used a dataset containing 5-10 trials each before and after tarsal amputation for each fly. The amputated legs technically did not have a stance phase, but they showed oscillations resembling steps. So we refer to the highest tarsal position during each oscillation as swing peak here for simplicity. We detected and manually corrected the timing of swing peaks of all steps for intact legs or all oscillations for amputated legs. Using this information, we could calculate the averaged frequency of steps or oscillations for each leg in each fly across multiple trials, which represent single data points in Fig. 1j. The step frequency in Extended Data Fig. 5c and Extended Data Fig. 7c was quantified in the same way.

For Fig. 5e and Extended Data Fig. 6d, we used a dataset that contained a single trial for each fly, in which the tracking of coxa-trochanter and femur-tibia joint angles in hind legs had minimal errors. Given many of the flies with LIN156 activated showed re-extension of tibia following initial flexion, we quantified the maximal flexion in 1 s after optogenetic stimuli onset before such re-extension.

For Fig. 5f,g we used 50-ms light pulses spaced by at least 1 s to activate the neuron. We aligned 105 trials from 3 flies by the stimulus onset and overlaid the femur-tibia angle to plot Fig. 5f. We then divided all the trials by whether re-extension of tibia followed the initial flexion and plotted the minimal femur-tibia joint angle following each stimulus to generate Fig. 5g.

For Fig. 6e, we manually scored whether a leg showed repeated stepping under 5-s red light stimuli for each leg of the flies in this dataset. For Fig. 6f, we used 5-ms light pulses spaced by at least 1 s to activate the neuron. We aligned 100 trials from 3 flies by the stimulus onset and calculated the average of each joint angle from all trials. We took the averaged joint angles during the first 0.5 s after stimulus onset to calculate the Z-score (number of standard deviations from the mean). For Fig. 6g, we manually scored every frame as either stance or swing for each leg. We can use this information to calculate the co-swing index to plot Fig. 6h. The co-swing index was defined by the total co-swing time of two limbs divided by the total time either of the two limbs were in swing during the optogenetic stimulation. This dataset included one video with 5-s stimuli for each of the 5 flies.

For Fig. 7d-g and Extended Data Fig. 7a,b, we used a dataset containing the same number of control trials and experimental trials for each fly, following the protocol illustrated in Fig. 7a. We detected and manually corrected all swing peaks (see above) for hindlegs. We then aligned all steps that happened during the specified time windows (10-s window delayed by 10 s after red light onset for Fig. 7d-g and 5-s windows as illustrated in Extended Data Fig. 7a,b) for each genotype and experimental condition using the swing peaks. We used a 300-frame (1.5 s) time window and placed the swing peak at frame 40 (200 ms before the swing peak was shown) to plot Fig. 7d. We truncated steps at boundaries of the above mentioned time windows or 8 frames (40 ms) before the next swing peak if the next step arose within the 1.5 s time window. For Fig. 7e-g, we calculated the mean of relevant parameters in all steps for each fly and condition. For Fig. 7e, we applied the Matlab function “fit” with input parameter “poly1” on the declining phase (from the maximal femur-tibia angle position to the boundary of the 1.5 s window, i.e., 1.3 s after the swing peak) of averaged femur-tibia angles in all steps for each fly and condition. We used the linearly fitted slope as a quantification of tibia flexion speed. For Fig. 7f, we calculated the averaged minimal femur-tibia joint angle shortly before each swing peak of all steps for each fly and condition. For Fig. 7g, we took the reciprocal of the average duration (time between two neighbouring swing peaks) of all steps for each fly and condition to calculate the step frequency.

#### Statistical analysis

Details about statistical tests and sample sizes are indicated in the Figure or Figure legends. All statistical analysis were performed in GraphPad Prism 8.0.

